# A computational workflow for assessing drug effects on temporal signaling dynamics reveals robustness in stimulus-specific NFκB signaling

**DOI:** 10.1101/2025.03.31.645599

**Authors:** Emily R. Bozich, Xiaolu Guo, Jennifer L. Wilson, Alexander Hoffmann

## Abstract

Single-cell studies of signal transduction have revealed complex temporal dynamics that determine downstream biological function. For example, the stimulus-specific dynamics of the transcription factor NFκB specify stimulus-specific gene expression programs, and loss of specificity leads to disease. Thus, it is intriguing to consider drugs that may restore signaling specificity in disease contexts, or reduce activity but maintain signaling specificity to avoid unwanted side effects. However, while steady-state dose-response relationships have been the focus of pharmacological studies, there are no established methods for quantifying drug impact on stimulus-response signaling dynamics. Here we evaluated how drug treatments affect the stimulus-specificity of NFκB activation dynamics and its ability to accurately code ligand identity and dose. Specifically, we simulated the dynamic NFκB trajectories in response to 15 stimuli representing various immune threats under treatment of 10 representative drugs across 20 dosage levels. To quantify the effects on coding capacity, we introduced a Stimulus Response Specificity (SRS) score and a stimulus confusion score. We constructed stimulus confusion maps by employing epsilon network clustering in the trajectory space and in various dimensionally reduced spaces: canonical polyadic decomposition (CPD), functional principal component analysis (fPCA), and NFκB signaling codons (i.e. established, informative dynamic features). Our results indicated that the SRS score and the stimulus confusion map based on signaling codons are best-suited to quantify stimulus-specific NFκB dynamics confusion under pharmacological perturbations. Using these tools we found that temporal coding capacity of the NFκB signaling network is generally robust to a variety of pharmacological perturbations, thereby enabling the targeting of stimulus-specific dynamics without causing broad side-effects.

## INTRODUCTION

Signaling pathways respond to environmental stimuli and direct downstream gene expression. Environmental information may be encoded in the dynamics of signaling activity, constituting a temporal signaling code (1–3). In recent years, advances in live-cell imaging techniques and the optimization of analytical software have enabled the single-cell study of the temporal profiles of key signaling pathways, including NFκB (4–8), MAPKs (ERK, JNK, p38) (9,10), p53 (11), and their combinations (12,13). Temporal coding in these pathways has been shown to regulate gene expression responses and thereby the resulting biological functions, while dysregulation of this temporal coding is associated with various pathological conditions (1,14–16).

Theoretical studies have explored that the dynamics of signaling pathways present new opportunities for therapeutic intervention (16). Stimulus-specific signaling dynamics have been proposed as drug targets, as demonstrated in studies of NFκB (16), p53 (11), ERK dynamics (17,18), PI3K (19), and the MAPK signaling network (20). This emerging shift toward designing pharmacological interventions that target the dynamics of signaling offers greater precision than traditional approaches targeting steady-state dose-response relationships.

While the theoretical work established that pharmacological perturbation signaling dynamics is possible, it remains unclear how stimulus-specific the effects of such perturbations are. Previous studies have primarily focused on specific biochemical reaction rates within abstracted and simplified regulatory networks to alter signal transduction specificity and fidelity (16,21,22). However, it remains unclear how these perturbations affect stimulus-specific responses by more complex, physiologically relevant signaling networks that are functionally pleiotropic. In fact, we currently lack established methods for evaluating the efficacy and specificity of pharmacological targeting stimulus-response signaling dynamics. To address these questions, we have selected a biological system that exhibits stimulus-specific signaling dynamics in response to a wide range of different stimuli and for which a mathematical model that accounts for the stimulus-response signaling dynamics is available: the NFκB signaling system in macrophages.

NFκB plays critical roles in regulating immune responses to pathogen associated molecular patterns (PAMPs), damage-associated molecular patterns (DAMPs), and cytokines via regulating transcription, cytokine production, cell division and death, etc (23). NFκB signaling dynamics encode information about immune threats (4,15,24,25) and regulate immune genes to achieve appropriate immune responses corresponding to the specific stimuli (26– 33). Loss of specificity in immune cell responses has been associated with autoimmune diseases (4), and other pathological conditions (15,34). Recently, we identified six informative dynamic features of NFκB signaling, termed “signaling codons”, that together are sufficient to encode stimulus-specific information (4). Specifically, “Speed” (Speed) captures the activation speed, “Peak Amplitude” (Amp) refers to the highest response level, “Duration” (Dur) captures the total time NFκB levels remain above an activation threshold, “Total Activity” (AUC) measures the overall accumulation of NFκB activity, while “Early vs. Late” (EvL) represents the front-loading of NFκB activity and “Oscillatory Power” (Osc) quantifies the oscillation properties of the signaling trajectory.

Prior work also described a mathematical model of the NFκB signaling system that accounts for the temporal trajectories of NFκB that are representative of experimental single-cell stimulus-response datasets (4). The model consists of receptor-associated signaling modules (receptor module) that respond to five representative ligands (TNF representing cytokines, LPS, CpG, and Pam3CSK representing bacterial derived molecules, Poly(I:C) representing virus derived molecules) and a common core module that includes the kinases TAK1 and IKK and the IκBα-NFκB negative feedback loop. The topology of this mathematical model are well supported by decades of experimental studies and the parameterization is based on numerous publications starting in 2002 (26,35).

To assess the impact of pharmacological perturbations on the integrity of the NFκB signaling network’s temporal coding capacity (i.e. its ability to convey stimulus-specific responses), we selected 10 inhibitors targeting biomolecules involved in NFκB signaling pathways, each tested at 20 dosages. Leveraging the well-established mathematical model of the NFκB signaling network allowed us to generate synthetic data representing cellular responses to different stimuli under these drug regimes. With this immense amount of synthetic perturbation data, we developed computational methods—stimulus confusion maps and stimulus confusion scores—to evaluate alterations in stimulus-response specificity resulting from pharmacological perturbations of the NFκB signaling network. We demonstrated the capacity of stimulus confusion maps to (1) identify pharmacological perturbations that may enhance or diminish stimulus-specific NFκB signaling dynamics; (2) select drug regimens that maximize or minimize the stimulus specificity of NFκB pathways; and (3) evaluate the side effects of drugs on altering the stimulus-response specificity of NFκB temporal patterns. Applying these tools shows the robustness of temporal coding capacity of the NFκB signaling network under a variety of pharmacological perturbations.

## RESULTS

### A computational workflow to assess how stimulus-specific NFκB signaling dynamics may be modulated

Pharmacological perturbations can affect the temporal coding capacity of the innate response network that ultimately controls the dynamical activation of NFκB. To investigate this at scale, we developed a computational workflow. The workflow is anchored by an experimentally validated systems model comprised of interconnected ordinary differential equations (ODEs) (4). This model recapitulates the temporal trajectories of nuclear NFκB activity in response to a range of doses of each of five pro-inflammatory ligands, and shows a high degree of specificity in terms of ligand- and dose-specific NFκB dynamics and therefore high temporal coding capacity. The model comprises 94 biochemical reaction that are controlled by 126 kinetic parameters, which may be subject to pharmacologic perturbation. Thus, the model may be used to generate data that addresses how pharmacological perturbation may modulate the coding capacity of the innate immune signaling network (Figure 1A).

**Figure 1.**
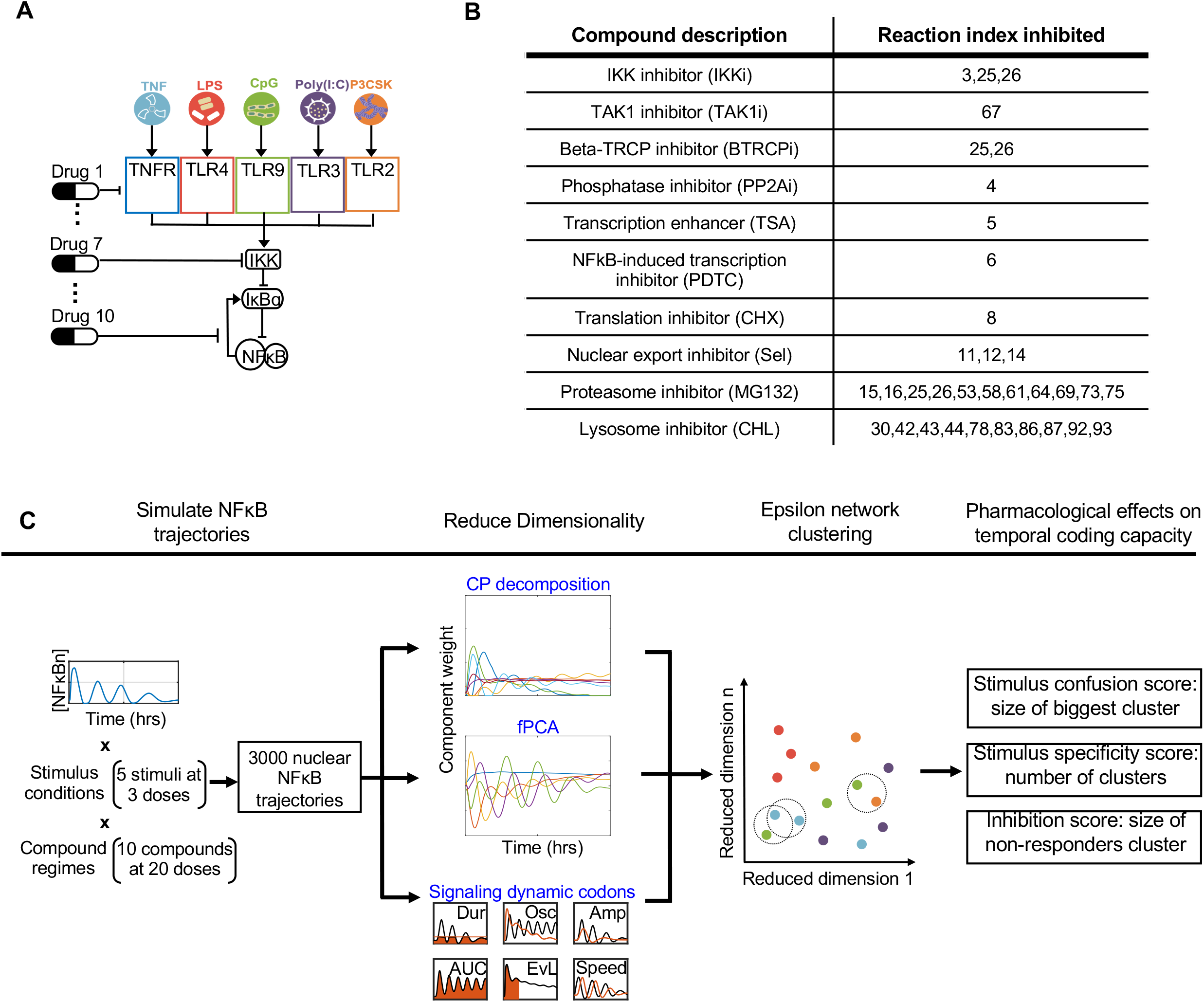
A computational workflow for identifying pharmacological perturbations that alter the temporal coding capacity of the NFκB signaling network. (A) Schematic of the NFκB network model. Five ligands bind to their cognate receptors triggering associated signaling modules that activate the kinase IKK which causes degradation of IκBα and hence de-repression of NFκB activity. There are numerous pharmacological agents that interfere with different reactions within this NFκB signaling network. (B) Table of pharmacological compounds that inhibit biochemical processes within the NFκB signaling network. Numbers indicate the kinetic parameters that are affected by the indicated compound. (C) The computational workflow begins with simulations that yield NFκB trajectories in response to 5 ligands (TNF, LPS, CpG, Poly(I:C), and Pam3CSK) at low/medium/high doses in the presence of pharmacological treatment with one of 10 compounds at one of 20 doses. The resulting 3000 trajectories are defined by 481 timepoints (every 5 min for 8 hrs) which are then reduced through (1) canonical polyadic (CP) decomposition, (2) functional principal component analysis (fPCA), or (3) quantifications of previously identified informative trajectory features, known as signaling codons. Using the outputted features from dimensionality reduction, epsilon network clustering is applied to the 15 stimulus conditions (colored by stimuli) within a drug regime. Cluster-based stimulus confusion, specificity, and inhibition scores are calculated to reveal the pharmacological effects on stimulus-response specificity of NFκB dynamics, also described as the temporal coding capacity of the NFκB signaling pathway.

We identified ten pharmacological compounds with well-described biochemical mechanisms of action (Figure 1B, Table S1). As such, we could map each compound to the affected kinetic parameters within the ODE model. This library includes compounds that impact the core module: IKK inhibitor (IKKi) (36), TAK1 inhibitor (TAK1i) (36), Beta-TRCP inhibitor (BTRCPi) (37,38), Phosphatase inhibitor (PP2Ai) (39,40), Transcription enhancer (TSA) (41), NFκB-induced transcription inhibitor (PDTC) (42), translation inhibitor (CHX) (42), and nuclear export inhibitor (Sel) (43). A proteasome inhibitor, MG132, inhibits the degradation of IκBα, whether complexed to NFκB or not, in the core module, and several receptors and their associated complexes in receptor modules (42). Lysosome inhibitor (CHL) inhibits the degradation of CD14 and several toll-like receptors (TLRs) trafficking to the plasma membrane and endosomes (44).

In order to quantify the stimulus-specific temporal coding capacity of NFκB under pharmacological perturbation, we designed a computational workflow. The workflow consists of the following steps: (1) generation of 3000 NFκB simulated trajectories in response to 15 stimulus conditions and under 200 pharmacological alterations (10 drugs at 20 doses resulting in 200 drug doses, DD) to our ODE model; (2) dimensionality reduction of the simulated trajectories using either matrix-based decomposition methods or decomposition into informative dynamic signaling features that were identified using information theory (4); in the reduced dimensional space, epsilon network clustering of simulated trajectories; (4) quantification of the confusion and specificity effects within each regime through the calculation of corresponding scores (Figure 1C). The goal of this workflow is to characterize the extent to which drug treatments (ranging from attenuating and enhancing effects on NFκB activation) may affect the stimulus-specific dynamics of the NFκB signaling module, either through maintaining or diminishing its temporal coding capacity.

### Generation of 3000 pharmacologically perturbed NFκB temporal trajectories

Using the NFκB systems model, we first generated stimulus-responsive nuclear NFκB temporal trajectories in response to 5 ligands (TNF, LPS, CpG, Poly(I:C), and Pam3CSK) at 3 doses (15 total stimulus conditions). These 15 trajectories are stimulus-specific and hence encapsulate NFκB’s temporal coding capacity. Next, we applied the 10 previously described pharmacological compounds to our model by modifying their corresponding kinetic parameters at 20 log-linearly spaced drug doses (DD) ranging from 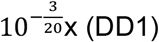 to 10^−3^x (DD20). This resulted in 3000 simulated NFκB trajectories, representing nuclear NFκB concentration over an 8-hour stimulation time course (Figure 2).

**Figure 2.**
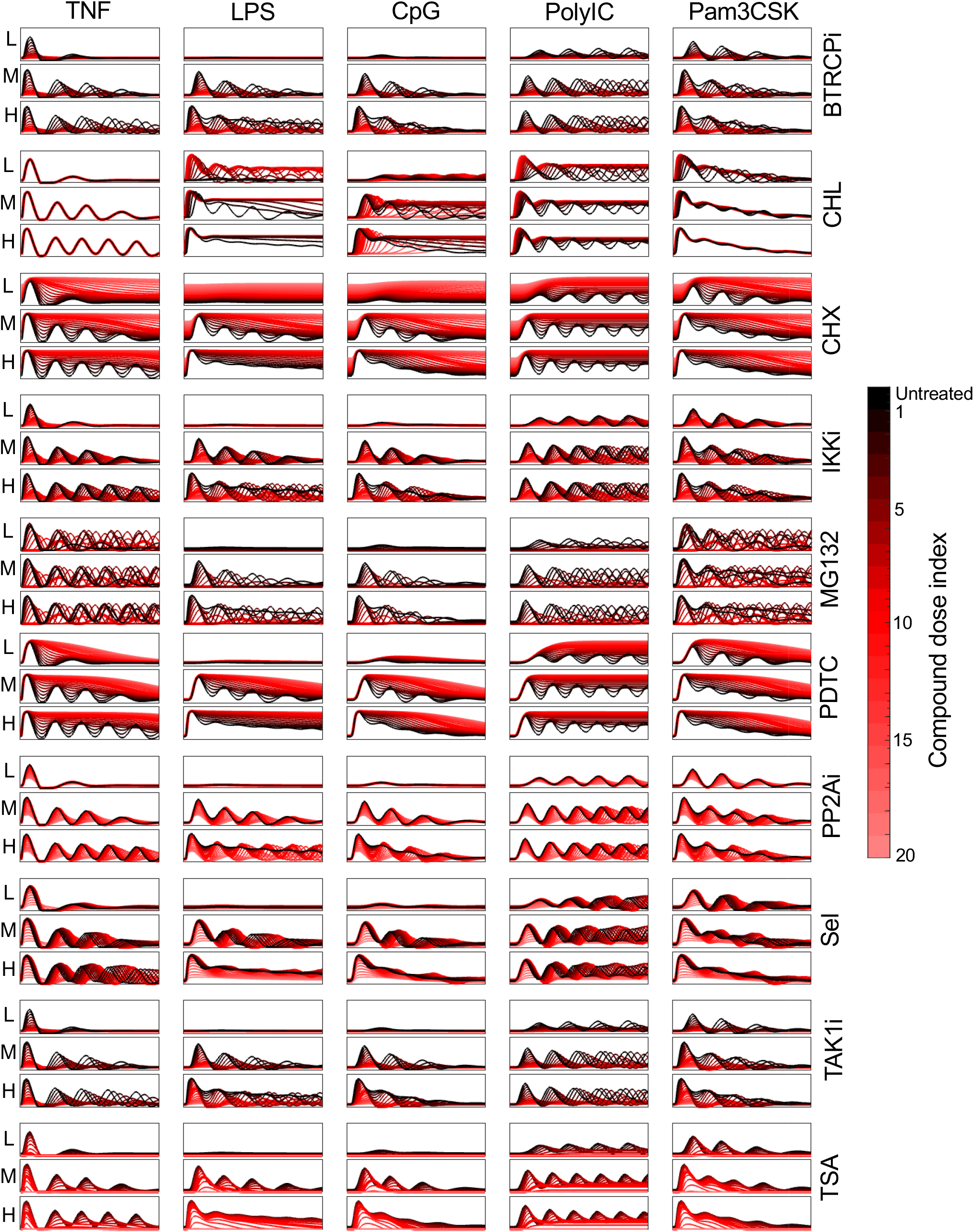
NFκB signaling trajectories under pharmacological perturbations. Stimulus-specific NFκB trajectories (0 – 8 hours) following computational compound perturbations. The stimulating ligands are indicated at the top labels. The doses of these stimuli (Low, Medium, and High) are denoted by small labels on the left. A total of 21 trajectories are presented in each graph, from the untreated condition (depicted in black) to a compound dose (CD) of 20 (depicted in bright red). The colorbar on the right corresponds to specific compound dose indices.

Visual inspection of the simulated trajectories suggests that across all 15 stimulus conditions, most compounds cause a reduction in NFκB activation with increasing compound doses. Exceptions include the transcription and translation inhibitors PDTC and CHX and the lysosome inhibitor CHL. By inhibiting the production of the NFκB inhibitor, IκBα, PDTC and CHX treatment enhances nuclear NFκB activity. Similarly, with increasing doses of CHL, lysosome-dependent receptor degradation decreases, resulting in maintained (TNF-stimulated) and enhanced (LPS, CpG, Poly(I:C), and Pam3CSK-stimulated) activation of the NFκB signaling network.

However, it remains unclear how the stimulus-specific temporal coding capacity of NFκB dynamics is affected, especially in conditions of attenuating drug treatments. For example, while Sel and MG132 both reduce NFκB activation at increasing doses, they do so through altering signaling dynamic features that could thereby affect the stimulus-specificity of the stimulus-response trajectories. Across stimuli, we observe that at intermediate drug doses, Sel generally maintains the same trajectory shape as each stimuli’s untreated trajectory whereas MG132 leads to modulation of oscillatory behavior. Thus, to elucidate compound-specific effects on stimulus-specific signaling dynamics, we applied the next step in our computational workflow, dimensionality reduction, in order to represent the 3000 complex temporal trajectories by key, interpretable features.

### Defining stimulus-response specificity landscapes for drug regimes

NFκB signaling is stimulus-response specific, and a loss of this stimulus-response specificity has been linked to autoimmune diseases, such as Sjogren’s Syndrome. To quantify the stimulus-response specificity, we sought a dimensionality reduction method that reduced data complexity but encoded stimulus-specific information of the signaling dynamics. Given the immense high dimensional and time-dependent information within our dataset, we set out to test the utility of three different dimensionality reduction methods which prioritize distinct characteristics within our data: canonical polyadic decomposition, functional principal component analysis, and signaling codons. Within each low dimensional space, the temporal trajectories are represented by a method-specific feature set. While these techniques were applied to all 200 drug regimes, we selected two representative regimes, Sel at DD15 and MG132 at DD6 (for simplicity, referred to as Sel and MG132 for the remainder of this section), to demonstrate the functionality of each dimensionality reduction method.

We first searched for general trends in the pharmacological effects within the original trajectory space. In doing so, we observed that Sel treatment, the nuclear export inhibitor (Figure 3A), reduces NFκB oscillations while generally maintaining each trajectory’s initial peak (untreated vs. treated in Figure 3B). We also noted this treatment’s similar effects on trajectories stimulated by medium doses of LPS and CpG (stimulation indices 5 and 8). In addition, MG132, the proteasome inhibitor (Figure 3C), causes stimulus- and dose-specific alterations to NFκB dynamics (treated vs. untreated in Figure 3D). For instance, all TNF- and Pam3CSK-stimulated trajectories have uniform oscillations (simulation indices 1-3 & 13-15) which can be attributed to their respective receptor’s (TNFR and TLR2) proteasome-mediated degradation. For these 6 stimulus conditions, treatment with MG132 results in competition between reduced receptor degradation and reduced IκBα degradation which yields similar trajectory shapes (simulation indices 1-3 & 13-15 in Figure 3D). These Sel- and MG132-treated trajectories were then compared to their feature vectors in the lower dimensional spaces (described in detail below) to determine the extent to which each technique recapitulates temporal dynamics.

**Figure 3.**
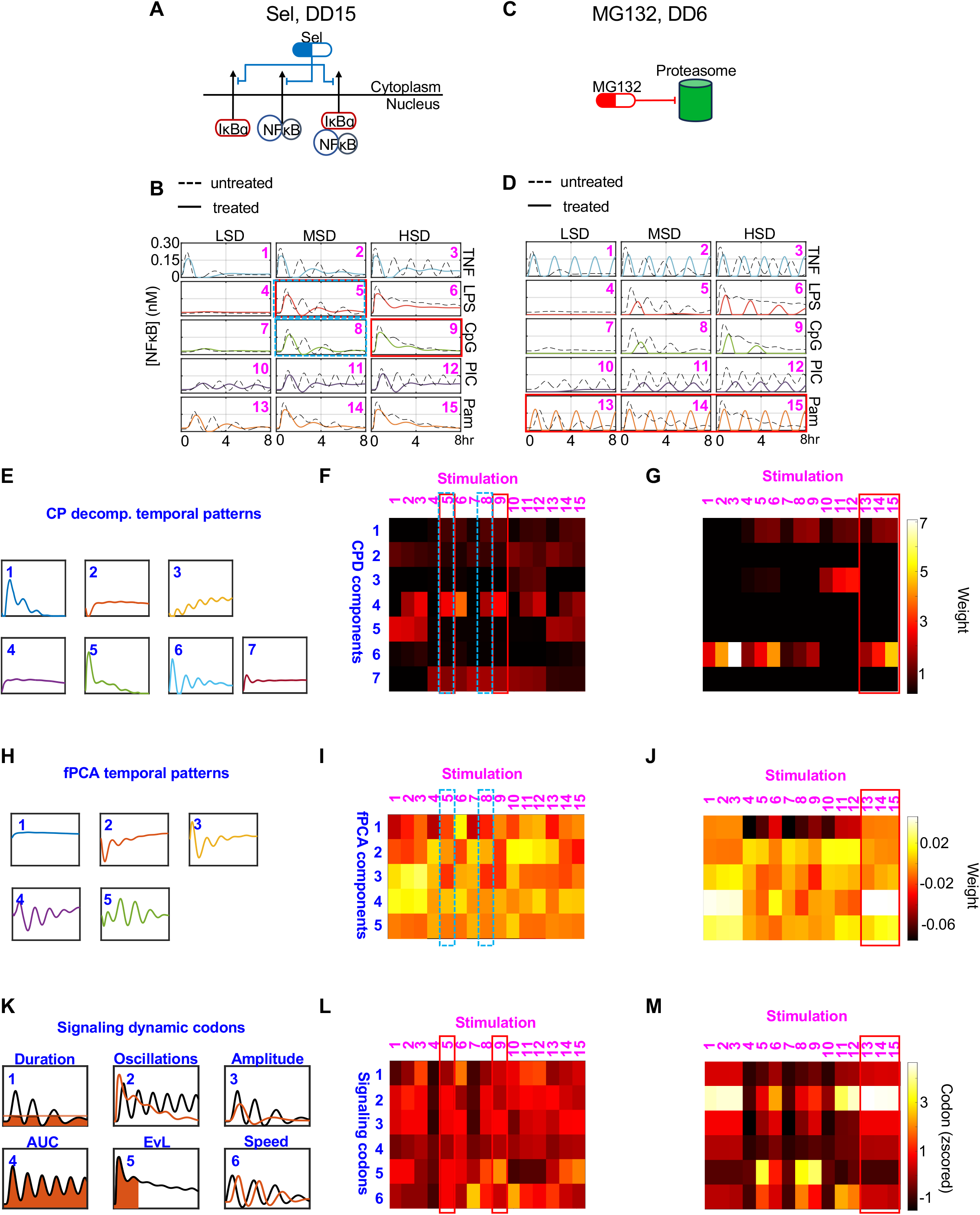
Feature/Reduced-dimensional representation of the trajectories. **(A)** Schematic of how example drug treatment Sel DD15 affects the NFκB signaling network. **(B)** Trajectories of NFκB activity over time for drug treatment Sel DD15 across 5 ligands and 3 doses. Solid colored lines represent the trajectories under drug treatment, while dashed lines depict untreated trajectories. Rows indicate different ligands (labeled on the right) and columns specify ligand doses (labeled on the top of the panel). Stimulation indices are labeled in the top right corner. **(C)** Schematic of how example drug treatment MG132 DD6 affects the NFκB signaling network. **(D)** Trajectories of NFκB activity over time for drug treatment MG132 DD6 shpwn as described in panel B. **(E)** Temporal patterns corresponding to seven CPD components identified for decomposing the entire NFκB trajectory dataset shown in Figure 2. **(F)** Heatmap representations of the weights of the indicated seven CPD components shown in panel E for the Sel DD15 data shown in panel B. **(G)** Heatmap representations of the weights of the indicated seven CPD components shown in panel E for the MG132 DD6 data shown in panel D. **(H)** Temporal patterns corresponding to five fPCA eigenfunctions components identified for decomposing the entire NFκB trajectory dataset shown in Figure 2. **(I)** Heatmap representations of the weights of the indicated five fPCA eigenfunctions shown in panel H for the Sel DD15 data shown in panel B. **(J)** Heatmap representations of the weights of the indicated five fPCA eigenfunctions shown in panel H for the MG132 DD6 data shown in panel D. **(K)** Schematic of the 6 dynamical features known as signaling codons, that were identified as informative of the stimulus-response NFκB trajectories. Temporal pattern and signaling codon indices are specified in the top left corner. **(L)** Heatmap representations of the normalized signaling codon values for the Sel DD15 data shown in panel B. **(M)** Heatmap representations of the normalized signaling codon values for the MG132 DD6 data shown in panel D.

The first dimensionality reduction approach is tensor decomposition (canonical polyadic decomposition, CPD), which provides a comprehensive factorized approximation of the data (45,46). The 3000 simulated trajectories were organized into a five-dimensional tensor (𝒜 ∈ 𝔽^481×5×3×10×20^), with dimensions representing ligand, ligand dose, drug, drug dose, and time. Employing non-negative CPD, we decomposed this tensor into 7 components, accounting for 87.5% of the variance observed in the original trajectory space (Figure S1A, Methods).

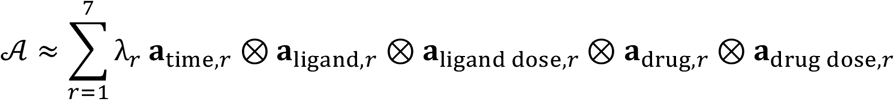

The factor weights along the time dimension (**a**_time,*r*_, *r* = 1, 2, …, 7) can be regarded as seven temporal patterns that encompass key dynamics within the original trajectory space (Figure 3E). Each trajectory is approximated as a weighted sum of these temporal patterns (Figure S1C, methods for detailed explanation).

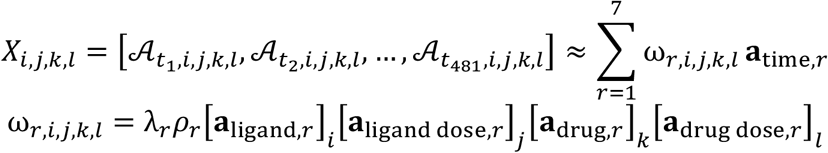

For each trajectory (X_*i,j,k,l*_), the 7 weights ({ω_*r,i,j,k,l*_}_*r*=1,2,…,7_) corresponding to the 7 temporal patterns represent its feature vector within the CPD-defined landscape. The component weights are calculated as the product of the component scalar, λ_*r*_, a scaling factor to correct for the inherent scale invariance of CPD, ρ_*r*_, and the factor weights in all but the time dimension (Figure S1B-D, see methods/supp for feature vector details). The landscapes (i.e., feature vectors) for the drug regimes highlighted above are visualized in Figure 3F-G. Focusing first on the Sel-treated landscape, we observed, for instance, that medium doses of LPS and CpG stimulation are mainly weighted in temporal patterns 4, and 7 (stimulation indices 5 and 8 in blue boxes of Figure 3F). However, these stimulus conditions have slightly different weights in temporal pattern 4, while in the original trajectory space medium doses of LPS and CpG stimulation are similar. Next, within the MG132-treated landscape, all doses of Pam3CSK stimulation are weighted in temporal patterns 1 and 6 to varying extents (stimulation indices 13-15 boxed in Figure 3G). This contradicts their nearly indistinguishable dynamics in the original trajectory space (stimulation indices 13-15 in Figure 3D). We speculate that these discrepancies in both regimes are because CPD disregards time-order information.

Considering the importance of temporal information, we also applied functional principal component analysis (fPCA) to our dataset. A major difference between fPCA and PCA is that the order of functional data carries continuous (i.e., temporal) information meaning that permutations of the multivariate data will lead to different results. We organized the simulated data into a 3000×481 matrix ([*X*_m,ti_]_*m*=1,2,…,3000;*i*=1,…,481_) containing the 3000 combinations of stimuli conditions (15) and drug regimes (200) at 481 timepoints. We found that the first 5 components account for 95.6% of the variance (Figure S2A, see methods for details). The resulting eigenfunctions characterize the dynamic temporal patterns (Figure 3H), and each trajectory can be approximated by the linear combination of these temporal patterns (Figure S2B-C).

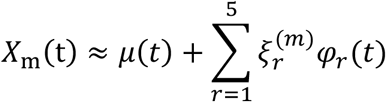

Like our CPD-defined feature vectors, the fPCA-defined feature vectors consist of fPCA scores 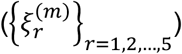 in the 5 components/temporal patterns (Figure 3H). The fPCA-defined Sel-treated landscape shows similar weights in the 5 temporal patterns for medium doses of LPS and CpG stimulation (stimulation indices 5 and 8 in blue dash boxes of Figure 3I). Regarding the MG132-treated landscape, while fPCA features better recapitulate the similarities between all Pam3CSK stimulated compared to CPD, fPCA still results in subtle variations in temporal patterns 2, 3, and 5 for these stimuli (stimulation indices 13-15 boxed in Figure 3J). Overall, this shows that encoding the timepoint order leads to a more meaningful representation of temporal trajectories.

Our final decomposition method leveraged the concept of NFκB signaling codons. The approach is to decomposes the trajectories into 6 informative dynamic features identified by an information maximizing algorithm that identified the dynamic features that best distinguished stimulus- and dose-specific NFκB responses (4). These 6 signaling codons include “Speed”, “Peak Amplitude”, “Duration”, “Area Under the Curve (AUC)”, “Early vs Late”, and “Oscillatory Power” (Figure 3K, see methods/supp for calculation details).

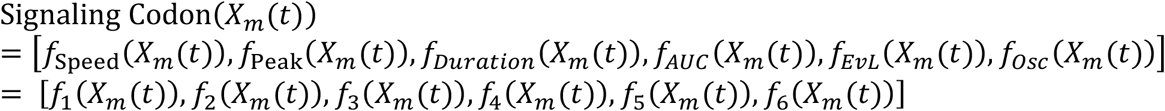

Normalized signaling codons values ({*f*_*s*_(*X*_*m*_(*t*)}_*s*=1,2,…,6_) make up the signaling codon-defined feature vectors used to derive the final drug regime landscapes (Figure 3L-M). In the signaling codon-defined Sel-treated landscape, we observe consistent, near average signaling codon values for medium doses of LPS and CpG stimulation (stimulation indices 5 and 9 in red boxes in Figure 3L). Interestingly, the subtle, yet significant discrepancies in the Oscillations and Early vs. Late codon for these conditions highlights the slightly more apparent three oscillatory peaks for the LPS-stimulated trajectory (stimulation indices 5 and 9 in red boxes in Figure 3L), which was not captured in the matrix-based decomposition landscapes. This underscores the importance of the capacity to discern both the number and timing of peaks in the signaling codon method. Further, the signaling codon-defined MG132-treated landscape also more adequately describes trajectory dynamics compared to the matrix-decomposition landscapes. This can be highlighted by the similar signaling codon values across Pam3CSK conditions, with an emphasis on their high oscillation power and relatively late NFκB activity (stimulation indices 13-15 highlighted/boxed in Figure 3M), whereas matrix methods decomposed these trajectories into weighted damped oscillation patterns.

In summary, based on the two drug regimes presented here, we found that matrix-based decompositions of temporal trajectories best recapture NFκB dynamics when the method emphasizes timeseries (fPCA) rather than structural information (CPD). Additionally, quantified signaling codons reflect a range of nuanced differences and general similarities between dynamic trajectories and may perform better in the analysis than the temporal patterns associated with matrix-based decompositions.

### Quantifying stimulus-response specificity of NFκB signaling pathway under drug regimes

To identify the extent to which a drug altered the signaling-specificity of NFκB, we clustered trajectories under the same drug treatment based on the similarity of their dynamic patterns (Figure 4A). We implemented epsilon network clustering on the original trajectories and in the previously described low dimensional feature spaces for five selected drug regimes (Figure S3A): CPD (using 7 components, 7C CPD, and extended to 40 components, 40C CPD), 5 component fPCA (5C fPCA), and 6 signaling codons (Figure 4A). If signal specificity and coding capacity were maintained, this would yield 15 distinct clusters unique to each stimulus, where fewer, and more intermingled clusters would indicate a loss signal specificity and loss of coding capacity. All clustering results were then visualized in stimulus confusion maps, where off-diagonal dark points indicate converging responder trajectories in response to two stimuli, that therefore constitutes a case of coding confusion. In contrast, red points on the diagonal indicate successfully inhibited trajectories (Figure 4B-C, Figure S3). To illustrate, for the drug treatment Sel DD15, the objective stimulus cluster map shows three confusion clusters: medium dose of LPS and CpG; medium and high doses of Poly(I:C); medium and high doses of Pam3CSK. In addition to the confusion clusters, the response to low dose LPS is “non-responder” and the other trajectories exhibit stimulus-specific dynamics (Figure 4B). Visually, the signaling codon-defined cluster maps most similarly resembled the objective clusters (Figure S3B) across all 5 validation drug regimes (Figure 4B, S3C).

**Figure 4.**
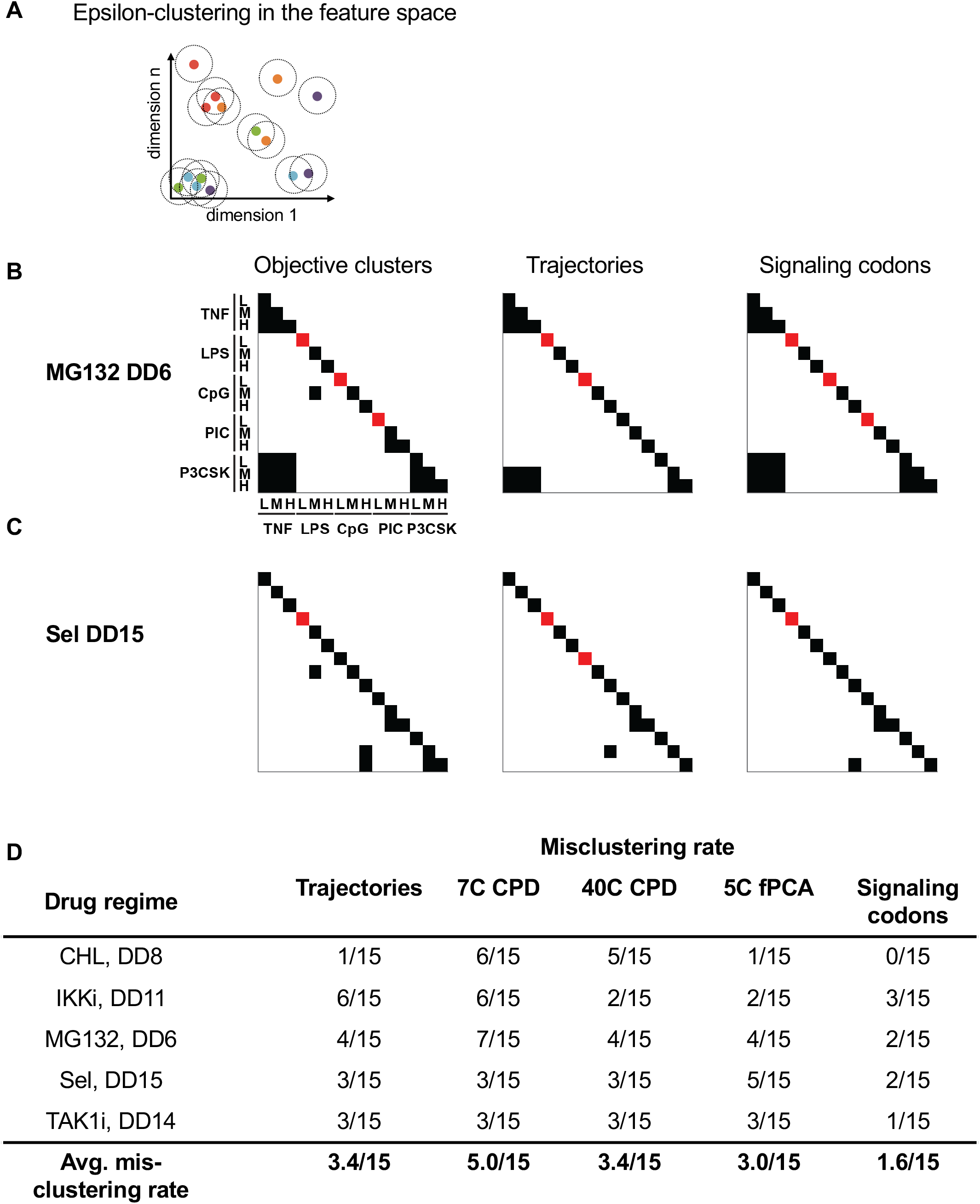
Construction of stimulus cluster maps for selected drug regimes. **(A)** Illustration of the definition of stimulus-response specificity (SRS) and stimulus-response confusion (SRC) scores *via* epsilon clustering of the high dimensional data. **(B)** Stimulus cluster maps constructed from epsilon network clustering results for drug treatment Sel DD15. Each row and column within one map correspond to a specific stimulus, as denoted on the left side of the left panel. Within each map, off-diagonal black squares represent responsive clusters and red squares on the diagonal represent inhibited NFκB signaling (non-responder). Left, middle, and right panels display ‘objective’ clusters, clusters derived from trajectory space, and clusters derived from signaling codon space, respectively. **(C)** Stimulus cluster maps constructed from epsilon network clustering results for drug treatment MG132 DD6. For labeling see panel B. **(D)** Misclustering rates (# of misclustered stimuli / 15 stimuli) for each feature space across the 5 random selected drug regime.

To quantitively determine which feature space gives the most reliable clusters, we calculated the misclustering rate under the optimal epsilon values in each feature space compared to the corresponding objective clustering (see methods for calculation details). Indeed, the signaling codon feature space had an average misclustering rate of 1.6/15 whereas most other feature spaces, including the original trajectory space, resulted in misclustering rates of more than 3.0/15 (Figure 4D). This shows that the low dimensional signaling codons prioritize key, dynamical features that may be deemphasized in the high dimensional trajectory space, ultimately resulting in a greater capacity to differentiate between stimulus-specific NFκB responses. Thus, we focused the remainder of our analysis to the signaling codon feature space.

### Quantifying pharmacological effects on the NFκB temporal coding capacity

The capacity to generate stimulus-specific responses may be defined as the temporal coding capacity of the NFκB signaling module. To quantify the signaling temporal coding capacity under each drug treatment, we introduced three metrics: the *stimulus-response specificity* (SRS), *confusion* (SRC), and *inhibition* (INH) *scores*. We defined these scores based on the reliable clusters obtained from epsilon-network clustering in the signaling codon feature space. Let S denote the set of the clusters of the responder trajectories under specific drug treatment:

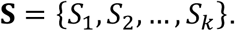

Then, the size of the largest cluster is defined as the *stimulus-response confusion score* (SRC), as all trajectories within one cluster are considered to be similar and potentially indistinguishable for NFκB target genes.

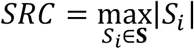

The *stimulus-response specificity score* (SRS) is the total number of clusters, which represents the number of uniquely identifiable NFκB signaling profiles under the same pharmacological perturbation.

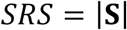

We also defined the *inhibition score* (INH) as the cluster size of non-responders which quantifies inhibition strength of each drug treatment.

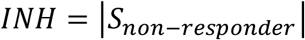

For all 200 drug regimes, we applied epsilon network clustering and used the resulting INH, SRC, and SRS scores to gain a comprehensive overview of trends in treatment effects on the temporal coding capacity of the innate NFκB signaling network. First, at high doses of treatment, we found that 5 compounds (e.g., BTRCPi and IKKi) cause complete inhibition while 2 compounds (CHL and CHX) lead to no inhibition of NFκB activity; 3 compounds (e.g., PP2Ai and Sel) selectively inhibit activity in a stimulus-dependent manner (Figure 5A). Additionally, the maximal stimulus confusion can be reached by compounds at intermediate (e.g., IKKi and MG132), high (e.g., CHX and PP2Ai), or relatively all (e.g., CHL) treatment doses (Figure 5B). With the network’s high innate temporal coding capacity (SRS = 13 for untreated condition), Sel is the only compound that enhanced temporal coding capacity (Sel at DD5-14 SRS = 15) (Figure 5C). Notably, regimes can modulate coding capacity by either partially inhibiting signaling while confusing all other stimuli (e.g., IKKi DD13 INH = 6, SRC = 9 Figure 5A,B) or by partially inhibiting signaling while still maintaining stimulus-specific responses (e.g., TAK1i DD12 INH = 10, SRS = 5, Figure 5A,C).Taken together, this suggest that confusion and inhibition can be selective. These distinct patterns of altering temporal coding capacity are summarized in more detail below.

**Figure 5.**
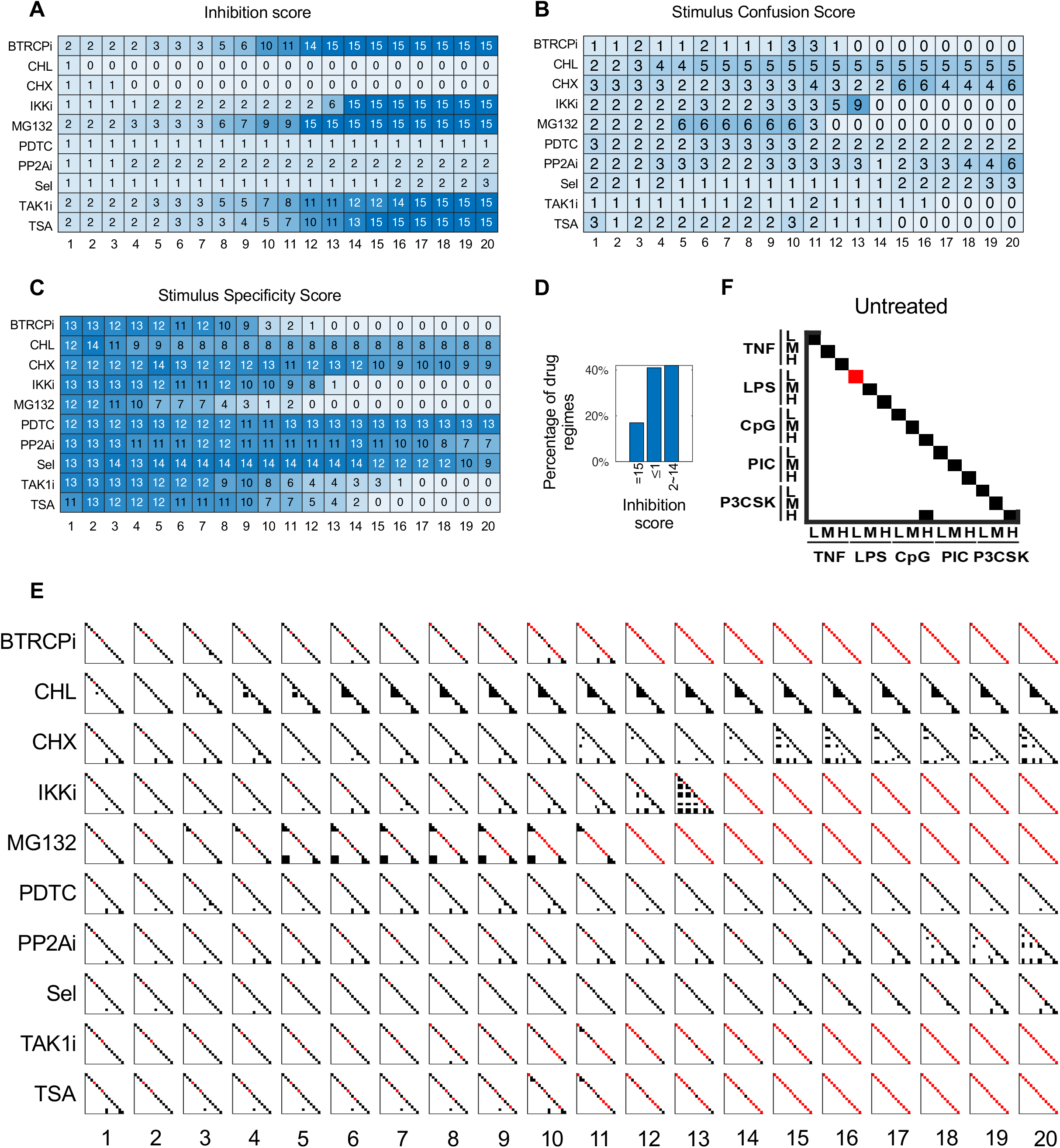
Signaling codon-derived quantification and visualization of the NFκB temporal coding capacity under each of 200 drug regimes. **(A)** Heatmap of inhibition scores for all 10 compounds across 20 compound doses treatments as indicated. **(B)** Heatmap of stimulus-response condusion (SRC) scores for all 10 compounds across 20 compound doses treatments as indicated. **(C)** Heatmap of stimulus-response specificity (SRS) scores for all 10 compounds across 20 compound doses treatments as indicated. **(D)** Bar plot of the percentage of drug regimes with specific range inhibition scores (specified in x-axis); corresponding to the three categories of drug regime functions. **(E)** Stimulus cluster map for the 15 stimulus conditions under no drug treatment (untreated). Each row and column within the map correspond to a specific stimulus, as denoted on the left and bottom side of the map. Off-diagonal black squares represent responsive clusters and red squares on the diagonal represent inhibited NFκB signaling (non-responder). **(F)** Stimulus confusion maps for all 10 compounds (far left labels) across 20 compound doses (bottom labels) treatments.

Using the previously described stimulus confusion maps, we next sought to investigate changes in temporal coding capacity at the level of specific stimulus conditions. For the 15 conditions under no drug treatment (i.e., untreated), 13 out of 15 were specific (SRS = 13), low dose LPS did not elicit a detectable NFκB response (INH = 1) and there were significant similarities between high doses of CpG and Pam3CSK-stimulated responses (SRC = 2) (Figure 5D).

Separately, the compound-treated stimulus confusion maps enabled us to find regimes that alter the temporal coding capacity of NFκB signaling pathway in three distinct ways: (i) complete inhibition of NFκB activity under all 15 stimulus conditions (Figure 5D, INH = 15) (ii-iii) partial stimulus-response specificity and confusion the rest of stimulus without (ii) (Figure 5D, INH <= 1) or with (iii) inhibition (Figure 5D, 1 < INH < 15). (i) High treatment doses of BTRCPi, IKKi, MG132, TAK1i, and TSA result in complete inhibition of NFκB activity (Figure 5A). As the compound dose increases, inhibition typically begins at low ligand doses and progresses to medium and high doses (Figure 5E). (ii) Drug regimes CHL DD20 and CHX DD20 exhibit partial stimulus-response specificity and partial confusion without inhibition (Figure 5A-C). CHL DD20 tends to confuse bacterial PAMPs such as LPS and CpG, while CHX DD20 primarily causes confusion between cytokines and PAMPs (Figure 5E). (iii) Compounds like MG132 DD6 and PP2Ai DD20 induce partial stimulus-response specificity and confusion, with the rest of stimuli inhibited. Under these regimes, the specificity scores are 7/15, with inhibition primarily affecting low-dose ligands and confusion occurring mostly at medium and high doses (Figure 5E).

The compound-treated stimulus confusion maps also allow for the identification of significant drug regimes that influence the NFκB signaling temporal coding capacity. For example, MG132 DD6 both induces confusion and inhibition for some stimulus conditions while maintaining specificity for the rest (Figure 5E). In the untreated regime, CpG and Pam3CSK are indistinguishable, whereas other stimuli, such as TNF and Pam, are distinct (Figure 5F). However, under MG132 DD6 treatment, the stimulus confusion map revealed that TNF and Pam become confused, while CPG and Pam become distinct (Figure S5A). By comparing trajectories and the corresponding signaling codon from untreated to treated regimes, we observed that TNF and Pam blur together, whereas Pam and CpG become clearly differentiated (Figure S5B). These results confirm the efficacy of signaling-codon-derived stimulus confusion maps in quantitatively estimating the effects of drug regime on altering NFκB temporal coding capacity.

### NFκB signaling temporal coding capacity is robust

While exploring pharmacological perturbations aimed at altering specific aspects of NFκB signaling, a possible side-effect is increased confusion among stimuli and conditions, thereby reducing the signaling temporal coding capacity. Thus, given the diverse pharmacological perturbations explored throughout this study, it is important to evaluate the robustness of the stimulus-specific temporal coding capacity of NFκB signaling pathway. we considered two key categories of robustness evaluation: (1) the degree to which stimulus specificity is maintained under pharmacological perturbation while altering specific NFκB dynamics; (2) the extent to which the drug-treated conditions are confused with the untreated conditions.

To broadly assess how much specificity is preserved within each drug regime, we computed the proportion of those that cause confusion for each pair of stimuli out of 200 drug regimes (Figure 6A). Stimuli pairs across cytokine and PAMPs are confused under few drug regime treatments. For instance, TNF vs. Poly(I:C) pair is confused by less than 10% of all drug regimes. Confusion among bacterial PAMPs occurs more frequently among drug regimes. This is exemplified by similar NFκB responses in the condition of high dose CpG and medium/high dose Pam3CSK under 89 drug regimes which can be explained by the similarity between their NFκB trajectories under the untreated condition.

**Figure 6.**
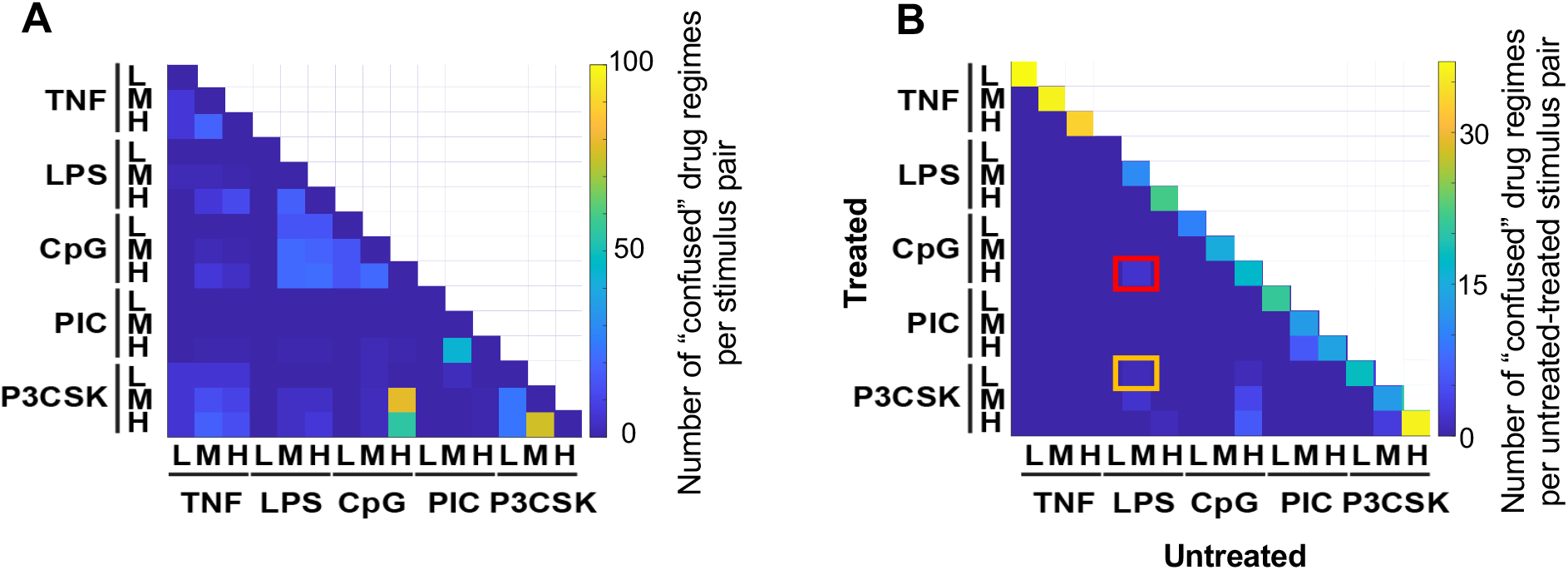
Evaluation of NFκB signaling network robustness. **(A)** Number of drug regimes where a given stimulus pair is “confused” within the same drug regime. Excludes regimes where both stimulated trajectories are completely inhibited by drug treatment. **(B)** Number of drug regimes where a given untreated-treated stimulus pair is “confused.” Columns represent the 15 untreated trajectories, rows represent all treated trajectories. Low dose LPS (column 4) induces an inhibitory response, resulting in no “confusion” across all treated conditions.

To evaluate the potential for confusion between drug-treated conditions and untreated conditions, we compared the pharmacologically perturbed conditions with the untreated NFκB signaling profile (Figure S6A-B). Most of the drug regimes showed minor confusions between treated and untreated conditions. The average frequency of same-stimuli confusion between treated and untreated conditions was 25/200, while across-stimuli confusion frequency was less than 10/200 (Figure 6B). Across-stimuli confusions between treated and untreated conditions were located primarily among the bacterial PAMPs. For example, Sel-DD13 treated medium-dose Pam3CSK stimulation is confused with untreated low-dose LPS stimulation, and IKKi-DD3 treated high-dose CpG stimulation resembles untreated medium-dose LPS stimulation (Figure S6C-D).

In summary, the pharmacological perturbations considered in this study have minor off-target effects of reducing NFκB signaling temporal coding capacity, and the temporal coding capacity of NFκB signaling remains robust against a variety of pharmacological perturbations.

## DISCUSSION

In this work, we introduced the stimulus-response specificity score and stimulus confusion maps to evaluate the efficacy and specificity of pharmacological perturbations in targeting stimulus-specific dynamical responses of the NFκB signaling pathway (Figure 4-5). We applied pharmacological perturbations to a well-established NFκB signaling network model, generating a large, simulated dataset of NFκB temporal trajectories (Figure 1-2). In generating low-dimensional representations of the temporal trajectories, we then found that signaling codons best recapitulated the stimulus- and perturbation-specific signaling dynamics, compared to traditional decomposition methods such as Canonical Polyadic Decomposition (CPD) and functional Principal Component Analysis (fPCA) (Figure 3-4). The application of the newly developed stimulus confusion maps to 200 drug regimens ultimately demonstrated the robustness of the temporal coding capacity of the NFκB signaling network (Figure 6).

One challenge in developing methods to evaluate the extent of pharmacological perturbation effects on stimulus-response specificity is the high dimensionality of time-series or trajectory data. To address this, we first reduced the time dimensions of the trajectories to a lower-dimensional representative space using CPD, fPCA, and signaling codon approaches (Figure 3). To assess the similarity or specificity of NFκB dynamics under drug treatment, we applied the epsilon network approach to the reduced-dimensional space, optimizing epsilon for each approach to generate confusion maps for each pharmacological perturbation. By comparing these confusion maps with objective clusters, we found that methods based on signaling codons provided the most accurate calculation of the stimulus confusion maps and the corresponding scores (Figure 4).

Why do signaling codons show superior performance over other methods of dimensionality reduction? Signaling codons are precise metrics that quantify informative dynamic features of NFκB signaling trajectories (4,24). They capture trajectory features that involve multiple time points and incorporate temporal order information. In contrast, decomposition approaches like canonical polyadic decomposition (CPD) and functional principal component analysis (fPCA) decompose trajectories into structural (CPD) or temporal (fPCA) components with corresponding weights (45,47). While these methods are effective in approximating or reconstructing the original trajectories through weighted combinations of components (45,47), they are less effective than signaling codons in quantifying the similarity and specificity of NFκB dynamics. CPD does not involve time point information and is insensitive to time-point order, while fPCA involves trajectories but the components are less precise and nimble than signaling codons, which are specific temporal features that were identified as being informative about the biological experimental signaling data.

By applying the newly developed stimulus confusion map to all 200 drug regimes, we found that the majority of the drugs inhibited NFκB signaling without losing the stimulus-specificity of the remaining NFκB stimulus responses, highlighting the robustness of the temporal coding capacity of NFκB signaling pathway (Figure 6). The confusion observed within select drug regimes, however, was primarily attributed to dose-related confusions within the same ligand stimulation rather than cross-ligand confusion (Figure 6). As immune response genes can decode dynamic features of NFκB signaling (26–31), this robustness in ligand distinction—even under drug treatment— might be indicative of minimal to no drug side effects or off-target effects. This could allow cellular NFκB responses in a specific condition to be diminished without causing confusion in immune responses to other stimuli. Such robustness of the temporal coding capacity of NFκB network to various pharmacological perturbations might be a result of evolutionary processes (48), as losing stimulus specificity could lead to pathological conditions (4,15). It will be of interest to characterize how the temporal coding capacity of other signaling modules such as ERK (extracellular signal-regulated kinase), Msn2 (yeast transcription factor), or p53 (tumor suppressor) (1), maybe be affected or remain robust in response to drug perturbations.

The described computational methods can be used to quantify the side effects of drugs targeting not only NFκB signaling, but other signaling systems as well, such as the p53 or ERK pathway (1), whose dynamics also encode stimulus information. To accomplish this, which dynamic features are informative about the stimulus and constitute signaling codons need to be determined for each signaling pathway and cell types (1). There are limitations to our workflow that might also present promising directions for future research. First, since single-cell data show high heterogeneity in signaling (5,49), incorporating cellular heterogeneity might provide new insights and present new challenges in the effective design of drugs. Second, in this study, we tested only idealized pharmacological perturbations, which are modeled to alter specific parameters within signaling network; the reality may be more complicated and how the drug effects are to be modelled must be refined to move the present study beyond the ‘proof-of-principle’ stage. Further, a broader drug library and whole-cell modeling may yield alternative insights regarding the robustness of the NFκB signaling pathway’s temporal coding capacity. Overall, our work presents an effective tool for evaluating to what extent pharmacological targeting may maintain stimulus-response specificity and reveals principles underlying the robustness of NFκB signaling dynamics.

## METHODS

### Simulation of NFκB trajectories under pharmacological perturbation

The previously developed 52-dimensional ordinary differential equation (ODE) model (4) was simulated. This model consists of five receptor modules that regulate the NFκB core signaling module. Drug-treated conditions were modeled by altering specific parameters as indicated in Figure 1B and Table S1. These changes involve multiplying certain parameters by values ranging from 10^−0.15^ to 10^−3^ in 20 linearly spaced steps, representing 20 different drug doses for each compound. Specifically, the IKK inhibitor (IKKi) reduces the values of parameters *k*_3_, *k*_25_, *k*_26_, while the TAK1 inhibitor (TAK1i) decreases parameter *k*_67_; the Beta-TRCP inhibitor (BTRCPi) reduces parameters *k*_25_ and *k*_26_; the phosphatase inhibitor (PP2Ai) modifies *k*_4_; the transcription enhancer (TSA) reduces *k*_5_; the NFκB-induced transcription inhibitor (PDTC) inhibits *k*_6_; the translation inhibitor (CHX) reduces *k*_8_; the nuclear export inhibitor (Sel) impacts parameters *k*_11_, *k*_12_, and *k*_14_; the proteasome inhibitor (MG132) reduces multiple parameters, including *k*_15_, *k*_16_, *k*_26_, *k*_26_, *k*_53_, *k*_58_, *k*_61_, *k*_64_, *k*_69_, *k*_73_, and *k*_75_; the lysosome inhibitor (CHL) inhibits parameters *k*_30_, *k*_42_, *k*_43_, *k*_44_, *k*_78_, *k*_83_, *k*_86_, *k*_87_, *k*_92_, and *k*_93_.

The ODE model was simulated using MATLAB’s ode15s solver. Simulations were conducted in two phases: an initial phase to establish a steady state under each condition (untreated and treated), followed by a second phase where different stimulation were applied. A total of 15 stimulations were applied, involving five ligands with three different doses each.

Specifically, the stimulation included 0.1 ng/mL, 1ng/mL, and 10ng/mL TNF; 1ng/mL, 10ng/mL, 100ng/mL LPS; 10nM, 100nM, 1000nM CpG; 1ug/mL, 10ug/mL, 100ug/mL Poly(I:C); and 10ng/mL, 100ng/mL, 1000ng/mL Pam3CSK. All simulation results were visualized using MATLAB.

### Dimensionality reduction of trajectory data

#### Canonical polyadic decomposition

Our simulated NFκB trajectories can be organized into a fifth-order tensor with dimensions of drug, drug doses, ligand, ligand doses, and time points. We turned to a tensor decomposition method, canonical polyadic decomposition (CPD), as our first dimensionality reduction technique. A key advantage of tensor decomposition over traditional techniques like principal component analysis is that it maintains the structural integrity of the data. We organized our simulated data into a fifth-order tensor, 𝒜 ∈ 𝔽^*i*×*J*×*K*×*L*×*T*^, where *I* represents the 5 ligands, *J* represents the 3 doses of each ligand, *K* represents the 10 drugs, *L* represents the 20 drug doses, and *T* represents the 481 timepoints. After applying CPD, 𝒜 was expressed as a linear combination of *R* rank-1 tensors (i.e., components):

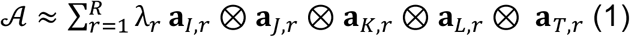

Here, λ_*r*_ is the component scalar, and **a**_*i,r*_, **a**_*J,r*_, **a**_*K,r*_, **a**_*L,r*_, and **a**_*T,r*_ are the factor vectors of the *r*^*th*^ component. The operator ⊗ denotes the outer product. The factor weights in the time dimension, **a**_*T,r*_, *r* = 1, 2, …, *R*, resemble distinct temporal patterns of NFκB trajectories under different stimulus conditions and drug treatment. We therefore referred to **a**_*T,r*_ as the key *R* temporal patterns within 𝒜.

For a specific trajectory stimulated by the *i*^*th*^ ligand at the *j*^*th*^ ligand dose and perturbed by the *k*_*th*_ drug at the *l*_*th*_ drug dose, 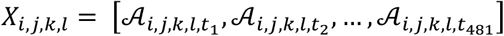, the linear combination of all temporal patterns is regarded as its approximated reconstruction:

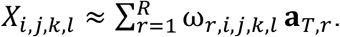

Each trajectory’s weight in the *r*^*th*^ temporal pattern/component (i.e., component weight), ω_*r,i,j,k,l*_, was calculated following Equation (2):

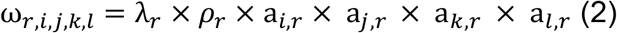

Due to the inherent scale invariance of CPD, where multiplying **a**_*i,r*_ by a constant and **a**_*T,r*_ by its reciprocal yields the same result, the solution to Equation (1) is not unique. To enable meaningful comparisons between components, we rescaled all temporal patterns to have an area of 20 and incorporated each pattern’s scaling factor ρ into the component weight calculations (Equation (2)).

Prior to decomposition, we normalized each element of 𝒜 within the drug dimension to ensure that the decomposition captured the effects of NFκB responses across the 10 drugs – each of which with a distinct mechanism of action – in a balanced manner. For drug treatment *k*, we normalized all timepoints as 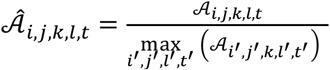.We applied CPD to 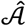 using the decomposition.non_negative_parafac function from the TensorLy package in python with ‘int’ = random and n_iter_max = 900 and all other settings left as the default parameters). Additionally, we tested ‘rank’ (i.e., number of components) = 1-8 to determine the optimal number of components. To quantify this, we first reconstructed the decomposition output, 𝒜_*reconstruct*_, using the cp_to_tensor function from the TensorLy package and then calculated the reconstruction error as 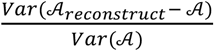 In doing so, we decided on *r* = 7 since higher values started to result in redundant temporal patterns. However, to test the limits of the method to accurately recapitulate and cluster timeseries data (see below), we also implemented it using 40 components.

Finally, for the *r* = 7 and *r =* 40 component decompositions, we organized each trajectory’s component weights into a feature vector, [ω_1,*i,j,k,l*_, ω_2,*i,j,k,l*_, …, ω_*r,i,j,k,l*_], respectively. These two separate sets of feature vectors represented the 7 component CPD- and 40 component CPD-defined landscapes.

#### Functional principal component analysis

Given that the simulated NFκB trajectories are time dependent, we employed functional principal component analysis (fPCA) as our second dimensionality reduction technique. The formalism of fPCA is similar to that of PCA with the key difference being that the data is decomposed into eigenfunctions rather than eigenvectors. Because of this, the variable (i.e., timepoint) order carries information, where random permutations result in different decompositions.

We first organized our data into a matrix, 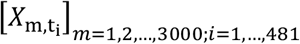 is the index representing a particular stimulus condition and a drug regime combination and *i* representing a given timepoint. Then, in applying fPCA to *X*, each individual trajectory, denoted by *X*_m_(*t*), was approximated according to Equation (3) where *µ*(*t*) represents the mean function used for centering *X* and 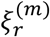 represents the m^th^ trajectory’s score (i.e., weight) in the r^th^ eigenfunction (i.e., functional principal component), *φ*_*r*_(*t*). As with PCA, *φ*_1_(*t*) captures the dominant pattern of variation within *X* with each subsequent component representing the dominant pattern of variation orthogonal to *φ*_1_(*t*), *φ*_2_(*t*), *φ*_3_(*t*), *φ*_*r*−1_(*t*). Therefore, each of the 3000 trajectories *X*_m=1,2,…,3000_(*t*) can be approximated by the linear combination of the eigenfunctions, each of which weighted by 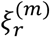.

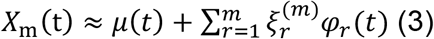

We implemented fPCA on our simulated trajectories using sckit-fda function in python. We opted to use a grid representation of *X* which we obtained with skfda.representation.grid.FDataGrid. With the resulting discretized object, we then initialized the fPCA class skfda.preprocessing.dim_reduction.FPCA using the default parameters ‘centering’ = True, ‘regularization’ = None, ‘component_basis’ = None, and ‘_weights’. To determine the optimal number of components (i.e., the minimum number that still captured a sufficient amount of variance in *X*), we varied the number of components parameter, ‘n_components,’ from 1-10. We then calculated and plotted the cumulative variance explained across all 10 fPCA decompositions. Setting ‘n_components’ = 5, we finally used the .fit_transform method to compute the first 5 components and their scores.

Similar to our interpretation of CPD weightings in the time dimension, we regarded the functional principal components as the key temporal patterns that dominated in our dataset. Therefore, for each trajectory *X*_m_, we organized their 5 weightings/scores in the first 5 functional principal components into a 5-dimensional feature vector. These vectors represented the fPCA-defined landscape.

#### Signaling codons

Six signaling codons are dynamic features capturing the stimulus-specific information encoded in macrophage NFκB trajectories (4). Specifically, “Speed” quantifies the activation response time, and “Peak” is the highest response activity. “Duration” measures the total time of NFκB level above a low threshold, while Area Under the Curve (“AUC”) quantifies the total integral of NFκB activity. Finally, Early Vs Late (“EvL”) represents the front-loading of NFκB activity, while Oscillatory Power (“Osc”) captures the oscillations due to IκBα negative feedback that are present in the trajectory. Below are the detailed definitions of each codon.

All the signaling codons are calculated using MATLAB. For an NFκB signaling trajectory, 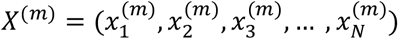 represents the simulated trajectory *m* across N time points with a time interval Δ*t*. 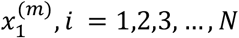, is the observation at time (*i* − 1) · Δ*t*.

1. Speed: for the trajectory *X*^(*m*)^, the local maxima peak sets λ_*peak*_ is defined as

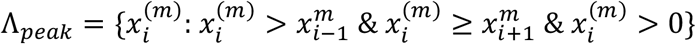

The index of the first peak time point is identified as:

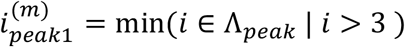

Speed is defined by:

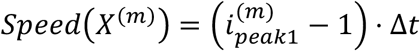
2. Peak: the maximal value of each trajectory:

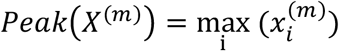
3. Duration:

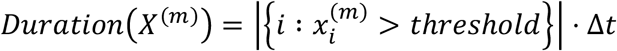
4. AUC:

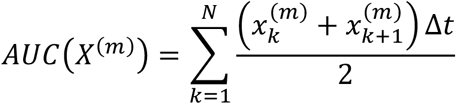
5. EvL:

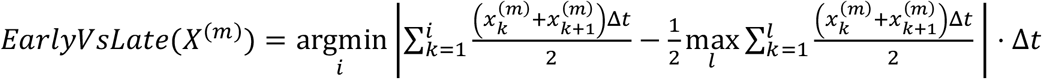
6. Osc:

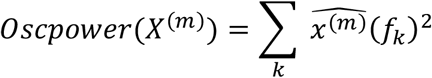

Where 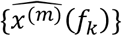 is the discrete Fourier transform of the trajectory 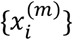, calculated using the fft function in MATLAB. 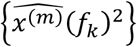 is the power spectral density. In the Osc formula, the summation is performed over the frequency range between 0.33 and 1 hour^-1^.

For each NFκB trajectory, the corresponding six signaling codons are calculated and saved in 6-dimensional vector. To obtain scores for each drug regime, these vectors are organized into a 15 × 6 matrix, with each row representing the NFκB trajectory under one of the fifteen stimulation conditions.

### Clustering of trajectories in original and dimensionality-reduced spaces

We developed an epsilon network clustering framework applied separately to five different feature spaces – 7 component CPD, 40 component CPD, 5 component fPCA, signaling codons, and the original trajectories – to quantify the similarities/differences between NFκB responses under distinct stimuli conditions for a given drug regime. To do this, we calculated the Euclidean distance between all pairwise feature vectors representing the 15 stimulus conditions in a drug regime within a given reduced dimensionality space. Smaller distances are indicative of a more similar response between two stimulus conditions. These distances were used as inputs for epsilon network clustering in which stimulus responses whose distances fell within a radius of ε were clustered together. Stimulus conditions are iteratively clustered according to ε, allowing for responses with a Euclidean distance ≥ ε to be clustered together if there is an intermediate condition with Euclidean distance ≤ ε for the aforementioned conditions.

To evaluate the accuracy of the generated clusters, we randomly selected five drug regimes and manually defined their corresponding objective clusters using independent individuals and taking their general consensus as the final set. For each of the original and 4 low dimensional feature spaces, the optimal ε, unique to each space, is defined by optimizing the ε that minimized the Misclustering Rate (MR) between epsilon network clustering and objective clusters, i.e.,

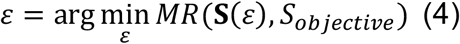

The misclustering rate between clusters are defined in the next paragraph. The optimized ε?were then applied to evaluate clustering performance within the different spaces.

To define MR between clusters A and B, *MR*(A, B), we first align the clusters from Partition A to those in Partition B in a way that minimizes mismatches. This defines the optimal mapping between the two partitions. We then identify mismatched elements and count them under the optimal mapping. Third, we calculate MR as the proportion of elements that are assigned to different clusters after optimal alignment.

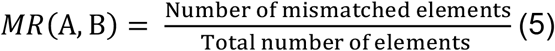

For example, for epsilon clustered results (Partition A of {1,2,3,…,15}) as

- Cluster A1: {1, 2, 3, 4, 5}
- Cluster A2: {6, 7, 8, 9, 10}
- Cluster A3: {11, 12, 13, 14, 15}

and objective cluster (Partition B)

- Cluster B1: {1, 2, 6, 7, 11}
- Cluster B2: {3, 4, 8, 9, 12}
- Cluster B3: {5, 10, 13, 14, 15}

The Optimal Mapping assignment is:

- A1 **↔** B1
- A2 **↔** B2
- A3 **↔** B3

Under this mapping, elements 3,4,5,6,7,10,11,12 – totaling eight elements – are mismatched. Thus the MR is

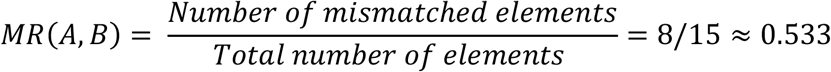

All above algorithms were implemented in MATLAB.

### Calculating stimulus-response specificity and confusion scores

We expanding our epsilon network clustering framework to all 200 drug regimes, using the predetermined optimal ε within each feature space. Clustering results were saved as confusion matrices and visualized in stimulus cluster maps. Each drug regime’s confusion matrix had a size of 15×15 with each column and row representing the different stimuli. Off-diagonal elements were assigned a value of 1 if the corresponding stimuli were in the same cluster and 0 otherwise. Non-responder stimuli for each drug regime were tracked using a separate vector. For each drug regime, since the confusion matrix is symmetric, only the lower triangular portion was visualized in stimulus cluster maps. Diagonal elements within the cluster maps, indicating the non-responder vector, were marked in red for inhibited trajectories classified as “non-responders.” For each stimulus pair, the number of drug regimes in which they were confused was tallied. All of the results were implemented and visualized in MATLAB.

For the five randomly selected drug regimes (Figure S3), non-responder trajectories are defined by an NFκB peak value below 0.05. Within the signaling codon space, we corrected the signaling codons EvL and Speed for non-responders to avoid non-sensical results.

When clustering, the non-responder cluster is defined as the one that overlaps the most with defined non-responder trajectories.

Because the signaling codon space achieved the smallest misclustering rate, we defined stimulus-specific and confusion scores solely within the signaling codon space. Given a drug regime’s set of clusters, **S** = {*S*_1_, *S*_2_, …, *S*_*k*_}, we considered stimuli that were within the same cluster and therefore elicited similar NFκB responses as confused while those within distinct clusters were defined as specific. For all drug regimes, non-responder NFκB trajectories were defined by an NFκB peak value below 0.05. These non-responder trajectories were contained in the non-responder cluster and used to quantify the Inhibition Score (INH):

*INH* = |*S_non-responder_*|. Among other responsive stimuli, the Stimulus Response Specificity (SRS) score was calculated as the total number of clusters: *SRS* = |**S**| while the Stimulus-Response Confusion (SRC) score was defined as the size of the largest cluster: 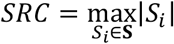. These calculations were implemented in MATLAB.

### Comparing drug-treated and untreated conditions

To compare drug-treated and untreated conditions, we constructed a confusion matrix in the signaling codon space by calculating the Euclidean distance between the untreated 15 stimulus-response vectors and their drug-treated counterparts. For responder conditions, distances less than the optimal ε were classified as confused and assigned a value of 1; otherwise, they were assigned 0. Non-responsive treated conditions were compared with the untreated low-dose LPS condition and marked in red. For each stimulus pair, the total number of drug regimes (out of 200) in which they were confused with untreated conditions was computed and visualized (Figure 6B). All the results were calculated and visualized in MATLAB.

## Code availability

All codes are available at GitHub (https://github.com/Xiaolu-Guo/Pharmacological_Perturbation_NFkB).

## Data availability

All data is deposited in Mendeley (https://data.mendeley.com/preview/t3dbj86sjx?a=f4fac707-ab61-49e4-a45b-b3bdebbe21cb).

## Author Contributions

A.H., X.G., and E.B. and designed the research. E.B. and X.G. developed the computational workflow, generated and analyzed all simulation data. X.G. and E.B. wrote and all authors edited the manuscript. A.H. and J.W. provided supervision and funding.

## Acknowledgements

The work is supported by National Institutes of Health (R01AI173214) to AH.

## References

1. Purvis JE, Lahav G. Encoding and Decoding Cellular Information through Signaling Dynamics. Cell. 2013 Feb 28;152(5):945–56.

2. Alexander RP, Kim PM, Emonet T, Gerstein MB. Understanding modularity in molecular networks requires dynamics. Sci Signal. 2009 Jul 28;2(81):pe44.

3. Behar M, Hoffmann A. Understanding the temporal codes of intra-cellular signals. Curr Opin Genet Dev. 2010 Dec;20(6):684–93.

4. Adelaja A, Taylor B, Sheu KM, Liu Y, Luecke S, Hoffmann A. Six distinct NFκB signaling codons convey discrete information to distinguish stimuli and enable appropriate macrophage responses. Immunity. 2021 May 11;54(5):916–930.e7.

5. Tay S, Hughey JJ, Lee TK, Lipniacki T, Quake SR, Covert MW. Single-cell NF-kappaB dynamics reveal digital activation and analogue information processing. Nature. 2010 Jul 8;466(7303):267–71.

6. Zhang Q, Gupta S, Schipper DL, Kowalczyk GJ, Mancini AE, Faeder JR, et al. NF-κB Dynamics Discriminate between TNF Doses in Single Cells. cels. 2017 Dec 27;5(6):638–645.e5.

7. Rahman SMT, Singh A, Lowe S, Aqdas M, Jiang K, Narayanan HV, et al. Co-imaging of RelA and c-Rel reveals features of NF-κB signaling for ligand discrimination. Cell Reports [Internet]. 2024 Mar 26 [cited 2025 Feb 23];43(3). Available from: https://www.cell.com/cell-reports/abstract/S2211-1247(24)00268-7

8. Selimkhanov J, Taylor B, Yao J, Pilko A, Albeck J, Hoffmann A, et al. Accurate information transmission through dynamic biochemical signaling networks. Science. 2014 Dec 12;346(6215):1370–3.

9. Davies AE, Pargett M, Siebert S, Gillies TE, Choi Y, Tobin SJ, et al. Systems-Level Properties of EGFR-RAS-ERK Signaling Amplify Local Signals to Generate Dynamic Gene Expression Heterogeneity. Cell Systems. 2020 Aug 26;11(2):161–175.e5.

10. Peterson AF, Ingram K, Huang EJ, Parksong J, McKenney C, Bever GS, et al. Systematic analysis of the MAPK signaling network reveals MAP3K-driven control of cell fate. Cell Systems. 2022 Nov 16;13(11):885–894.e4.

11. Paek AL, Liu JC, Loewer A, Forrester WC, Lahav G. Cell-to-Cell Variation in p53 Dynamics Leads to Fractional Killing. Cell. 2016 Apr 21;165(3):631–42.

12. Hanson RL, Batchelor E. Coordination of MAPK and p53 dynamics in the cellular responses to DNA damage and oxidative stress. Molecular Systems Biology. 2022 Dec;18(12):e11401.

13. Luecke S, Guo X, Sheu KM, Singh A, Lowe SC, Han M, et al. Dynamical and combinatorial coding by MAPK p38 and NFκB in the inflammatory response of macrophages. Molecular Systems Biology. 2024 Aug 2;20(8):898–932.

14. Hanahan D, Weinberg RA. Hallmarks of Cancer: The Next Generation. Cell. 2011 Mar 4;144(5):646–74.

15. Luecke S, Sheu KM, Hoffmann A. Stimulus-specific responses in innate immunity: Multilayered regulatory circuits. Immunity. 2021 Sep 14;54(9):1915–32.

16. Behar M, Barken D, Werner SL, Hoffmann A. The Dynamics of Signaling as a Pharmacological Target. Cell. 2013 Oct 10;155(2):10.1016/j.cell.2013.09.018.

17. Bugaj LJ, Sabnis AJ, Mitchell A, Garbarino JE, Toettcher JE, Bivona TG, et al. Cancer mutations and targeted drugs can disrupt dynamic signal encoding by the Ras-Erk pathway. Science. 2018 Aug 31;361(6405):eaao3048.

18. Goglia AG, Wilson MZ, Jena SG, Silbert J, Basta LP, Devenport D, et al. A Live-Cell Screen for Altered Erk Dynamics Reveals Principles of Proliferative Control. cels. 2020 Mar 25;10(3):240–253.e6.

19. Madsen RR, Vanhaesebroeck B. Cracking the context-specific PI3K signaling code. Science Signaling. 2020 Jan 7;13(613):eaay2940.

20. Mitchell A, Wei P, Lim WA. Oscillatory stress stimulation uncovers an Achilles’ heel of the yeast MAPK signaling network. Science. 2015 Dec 11;350(6266):1379–83.

21. Komarova NL, Zou X, Nie Q, Bardwell L. A theoretical framework for specificity in cell signaling. Mol Syst Biol. 2005 Oct 18;1:2005.0023.

22. Behar M, Dohlman HG, Elston TC. Kinetic insulation as an effective mechanism for achieving pathway specificity in intracellular signaling networks. Proc Natl Acad Sci U S A. 2007 Oct 9;104(41):16146–51.

23. Hayden MS, West AP, Ghosh S. NF-κB and the immune response. Oncogene. 2006 Oct;25(51):6758–80.

24. Aqdas M, Sung MH. NF-κB dynamics in the language of immune cells. Trends Immunol. 2023 Jan;44(1):32–43.

25. Singh A, Sen S, Iter M, Adelaja A, Luecke S, Guo X, et al. Stimulus-response signaling dynamics characterize macrophage polarization states. cels [Internet]. 2024 Jun 5 [cited 2024 Jun 6];0(0). Available from: https://www.cell.com/cell-systems/abstract/S2405-4712(24)00146-7

26. Hoffmann A, Levchenko A, Scott ML, Baltimore D. The IkappaB-NF-kappaB signaling module: temporal control and selective gene activation. Science. 2002 Nov 8;298(5596):1241–5.

27. Werner SL, Barken D, Hoffmann A. Stimulus specificity of gene expression programs determined by temporal control of IKK activity. Science. 2005 Sep 16;309(5742):1857–61.

28. Lee REC, Walker SR, Savery K, Frank DA, Gaudet S. Fold-change of nuclear NF-κB determines TNF-induced transcription in single cells. Mol Cell. 2014 Mar 20;53(6):867–79.

29. Sen S, Cheng Z, Sheu KM, Chen YH, Hoffmann A. Gene Regulatory Strategies that Decode the Duration of NFκB Dynamics Contribute to LPS-versus TNF-Specific Gene Expression. Cell Syst. 2020 Feb 26;10(2):169–182.e5.

30. Cheng QJ, Ohta S, Sheu KM, Spreafico R, Adelaja A, Taylor B, et al. NF-κB dynamics determine the stimulus specificity of epigenomic reprogramming in macrophages. Science. 2021 Jun 18;372(6548):1349–53.

31. Ando M, Magi S, Seki M, Suzuki Y, Kasukawa T, Lefaudeux D, et al. IκBα is required for full transcriptional induction of some NFκB-regulated genes in response to TNF in MCF-7 cells. NPJ Syst Biol Appl. 2021 Dec 1;7:42.

32. Sheu KM, Hoffmann A. Functional Hallmarks of Healthy Macrophage Responses: Their Regulatory Basis and Disease Relevance. Annu Rev Immunol. 2022 Apr 26;40:295–321.

33. Sheu K, Luecke S, Hoffmann A. Stimulus-specificity in the Responses of Immune Sentinel Cells. Curr Opin Syst Biol. 2019 Dec;18:53–61.

34. Murray PJ, Wynn TA. Protective and pathogenic functions of macrophage subsets. Nat Rev Immunol. 2011 Nov;11(11):723–37.

35. Basak S, Behar M, Hoffmann A. Lessons from mathematically modeling the NF-κB pathway. Immunol Rev. 2012 Mar;246(1):221–38.

36. Gupta SC, Sundaram C, Reuter S, Aggarwal BB. Inhibiting NF-κB activation by small molecules as a therapeutic strategy. Biochim Biophys Acta. 2010;1799(10–12):775–87.

37. Winston JT, Strack P, Beer-Romero P, Chu CY, Elledge SJ, Harper JW. The SCFbeta-TRCP-ubiquitin ligase complex associates specifically with phosphorylated destruction motifs in IkappaBalpha and beta-catenin and stimulates IkappaBalpha ubiquitination in vitro. Genes Dev. 1999 Feb 1;13(3):270–83.

38. Benary U, Wolf J. Controlling Nuclear NF-κB Dynamics by β-TrCP-Insights from a Computational Model. Biomedicines. 2019 May 27;7(2):40.

39. Witt J, Barisic S, Schumann E, Allgöwer F, Sawodny O, Sauter T, et al. Mechanism of PP2A-mediated IKK beta dephosphorylation: a systems biological approach. BMC Syst Biol. 2009 Jul 16;3:71.

40. Ho WS, Wang H, Maggio D, Kovach JS, Zhang Q, Song Q, et al. Pharmacologic inhibition of protein phosphatase-2A achieves durable immune-mediated antitumor activity when combined with PD-1 blockade. Nat Commun. 2018 May 29;9(1):2126.

41. Saccani S, Pantano S, Natoli G. Two waves of nuclear factor kappaB recruitment to target promoters. J Exp Med. 2001 Jun 18;193(12):1351–9.

42. Behar M, Hoffmann A. Tunable signal processing through a kinase control cycle: the IKK signaling node. Biophys J. 2013 Jul 2;105(1):231–41.

43. Kashyap T, Argueta C, Aboukameel A, Unger TJ, Klebanov B, Mohammad RM, et al. Selinexor, a Selective Inhibitor of Nuclear Export (SINE) compound, acts through NF-κB deactivation and combines with proteasome inhibitors to synergistically induce tumor cell death. Oncotarget. 2016 Nov 29;7(48):78883–95.

44. Newton PT, Vuppalapati KK, Bouderlique T, Chagin AS. Pharmacological inhibition of lysosomes activates the MTORC1 signaling pathway in chondrocytes in an autophagy-independent manner. Autophagy. 2015;11(9):1594–607.

45. Kolda TG, Bader BW. Tensor Decompositions and Applications. SIAM Rev. 2009 Aug 6;51(3):455–500.

46. Chin JL, Chan LC, Yeaman MR, Meyer AS. Tensor-based insights into systems immunity and infectious disease. Trends Immunol. 2023 May;44(5):329–32.

47. Ramsay J. Functional Data Analysis. In: Encyclopedia of Statistics in Behavioral Science [Internet]. John Wiley & Sons, Ltd; 2005 [cited 2024 Sep 27]. Available from: https://onlinelibrary.wiley.com/doi/abs/10.1002/0470013192.bsa239

48. Kitano H. Biological robustness. Nat Rev Genet. 2004 Nov;5(11):826–37.

49. Altschuler SJ, Wu LF. Cellular Heterogeneity: Do Differences Make a Difference? Cell. 2010 May 14;141(4):559–63.

