## Supplementary Information for "A computational workflow for assessing drug effects on temporal signaling dynamics reveals robustness in stimulus-specific NFκB signaling"

### Figure S1

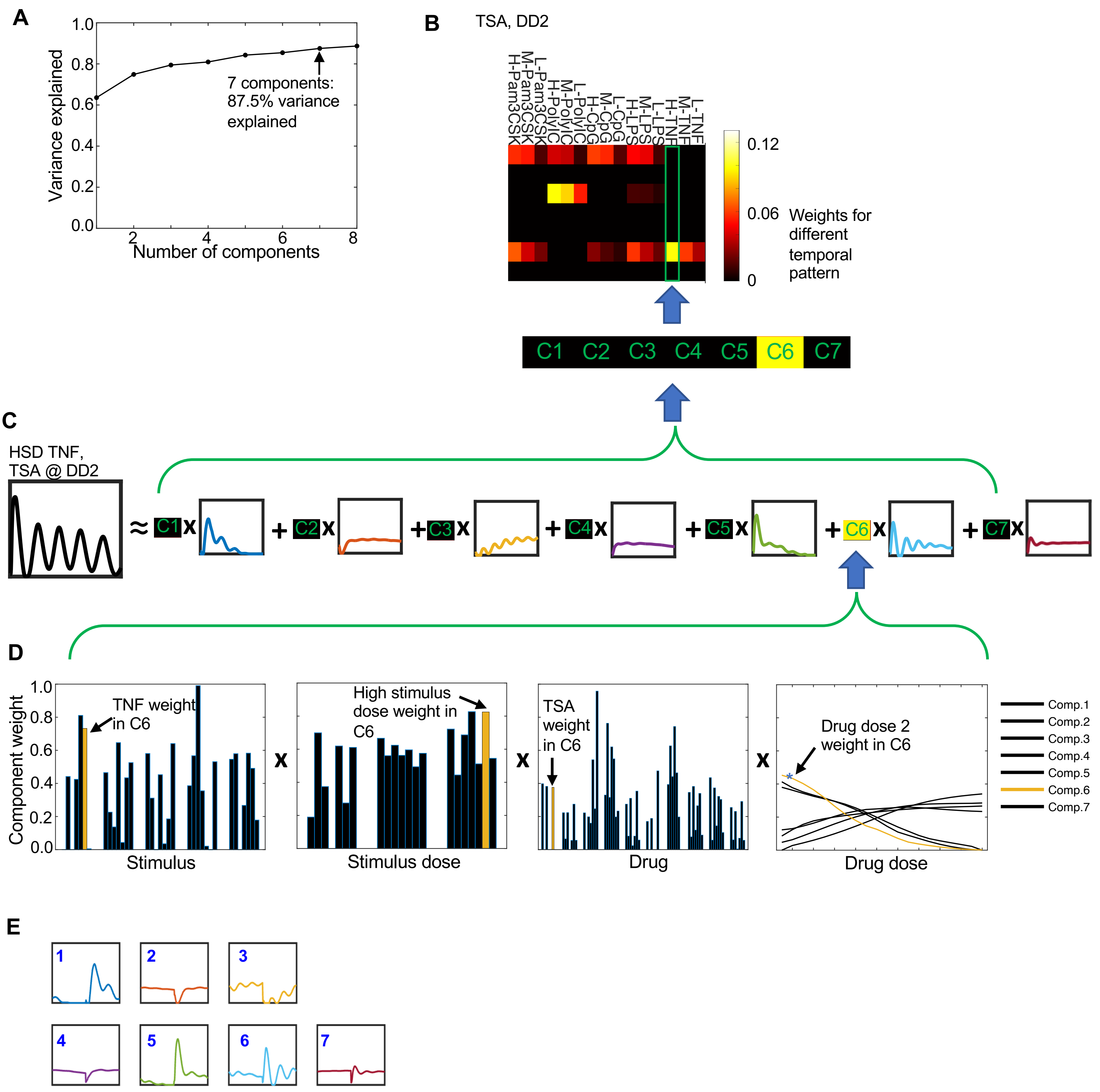
A
Variance explained
Number of components
7 components:
87.5% variance
explained
B
TSA, DD2
H-Pam3CSK
M-Pam3CSK
L-Pam3CSK
H-PolyIC
M-PolyIC
L-PolyIC
H-CpG
M-CpG
L-CpG
H-LPS
M-LPS
L-LPS
H-TNF
M-TNF
L-TNF
0.12
0.06
0
Weights for different temporal pattern
C1 C2 C3 C4 C5 C6 C7
C
HSD TNF,
TSA @ DD2
D
Component weight
TNF weight in C6
Stimulus
High stimulus dose weight in C6
Stimulus dose
TSA weight in C6
Drug
Drug dose 2 weight in C6
Drug dose
Comp.1
Comp.2
Comp.3
Comp.4
Comp.5
Comp.6
Comp.7
E

**Figure S1 Constructing landscape for drug regime from weights corresponding to seven components from the CP decomposition**

**(A)** R2X plot depicting the percent variance explained after the application of CP decomposition for 1 to 8 components.

**(B)** Heatmap of weights corresponding to seven temporal patterns signifying the drug TSA at dose 2 as an example defining the landscape for specific drug regime. Inside the heatmap, 15 rows depict 15 stimuli (5 ligands at 3 different doses). Colors within each row indicate the weights of the seven temporal patterns. These weights are the product of the component weights associated with the respective drug, drug dose, ligand, and ligand dose dimensions. The specific order of the 15 stimuli is outlined on the top of the figure, with L, M, and H denoting low-dose, medium-dose, and high-dose respectively. The weights of the temporal pattern (squares labeled by C1-C7 represent the weights for component 1-7 temporal patterns, respectively) stimulated by high-dose TNF under the TSA DD2 drug regime are specifically highlighted.

**(C)** An example of the high-dose TNF stimulated NFκB trajectory perturbed by TSA at drug dose index 2 is displayed on the left. This can be approximated by the weighted sum (weights indicated by colors in the squares) of the seven temporal patterns (displayed adjacent to the colored squares). The weights are the product of the component weights corresponding to the specific drug, drug dose, ligand, and ligand dose dimensions obtained from CP decomposition.

**(D)** The weights employed in the reconstruction of the simulated trajectory are the products of the weights in the dimensions of ligand, ligand dose, drug, and drug dose, with the yellow bars providing an example of calculating the weights for component 6 of (C). The subpanels and Figure 3E are the outcomes of Canonical Polyadic Decomposition (CPD) to nuclear NFκB time trajectory tensor, resulting in seven distinct components. These components are represented by their respective weights across various dimensions: Time (Figure 3E), Ligand, Ligand Doses, Drugs, and Drug Dose Index (this figure).

**(E)** Temporal patterns of CP decomposition components under time point order alterations. We altered the time point order of the original tensor by splitting it into two parts along the time dimension, and then taking their reversed concatenation (i.e., the altered-time-order tensor had a time point order of (241:481, 1:240). We then applied CP decomposition using 7 components for this altered tensor.

### Figure S2

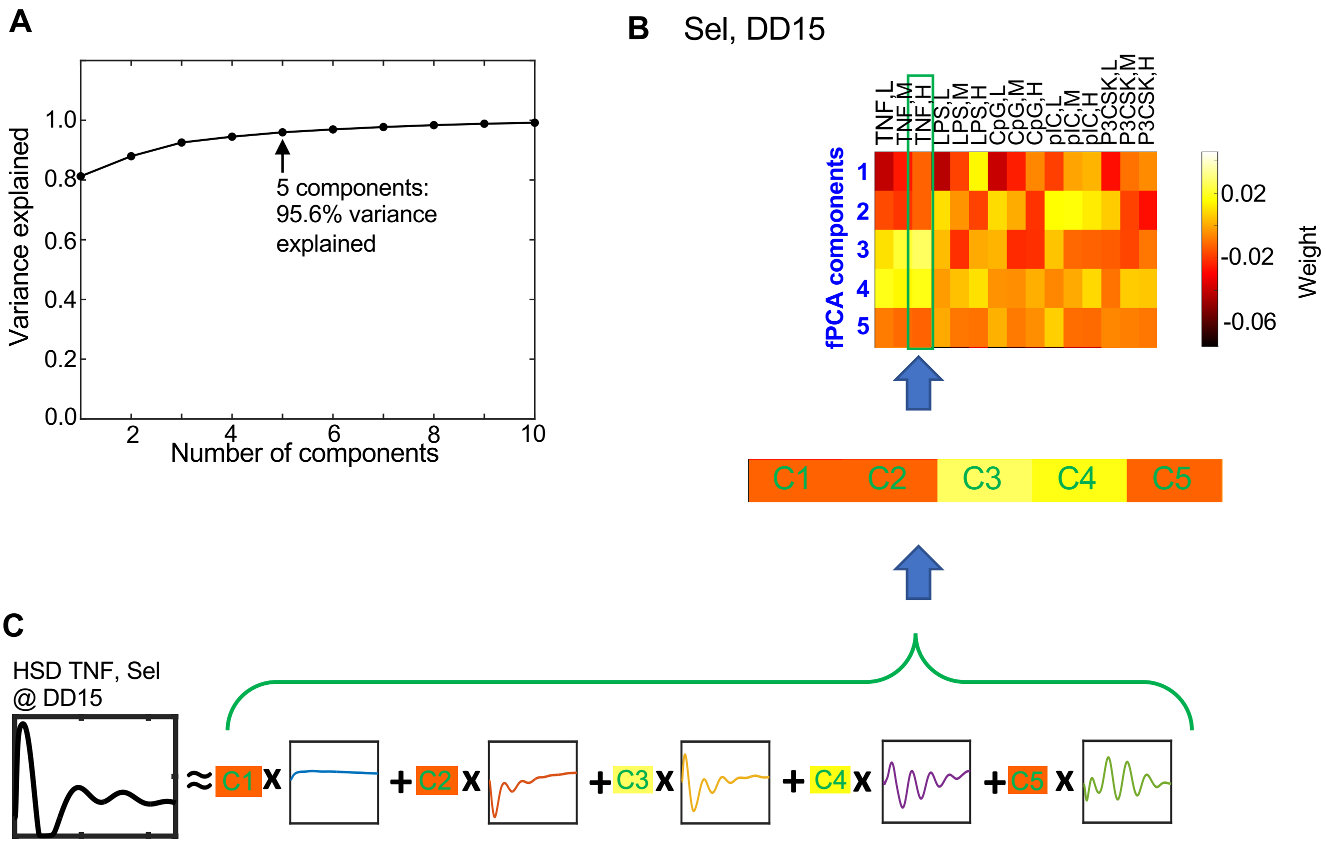


**Figure S2 Constructing landscape for drug regime from weights corresponding to five components from the fPCA**

**(A)** R2X plot depicting the percent variance explained after applying fPCA using 1-10 principal components.

**(B)** Heatmap of weights corresponding to five temporal patterns that are comprised in the drug regime Sel drug dose (DD) 15. The weights are the scores outputs from the decomposition. The 15 heatmap rows correspond to the 15 stimuli (5 ligands at 3 different doses) with their order annotated at the top of the heatmap (L = low-dose, M = medium-dose, H = high-dose). The squares labeled C1-C5 represent the five temporal pattern (i.e., components 1-5) weights for NFκB activity under high-dose TNF stimulation and perturbed by the Sel DD15 drug regime.

**(C)** An example of the high-dose TNF stimulated NFκB trajectory perturbed by Sel DD15 is displayed on the left. This can be approximated by the weighted sum (weights indicated by colors in the squares) of the five temporal patterns (displayed adjacent to the colored squares).

### Figure S3

A

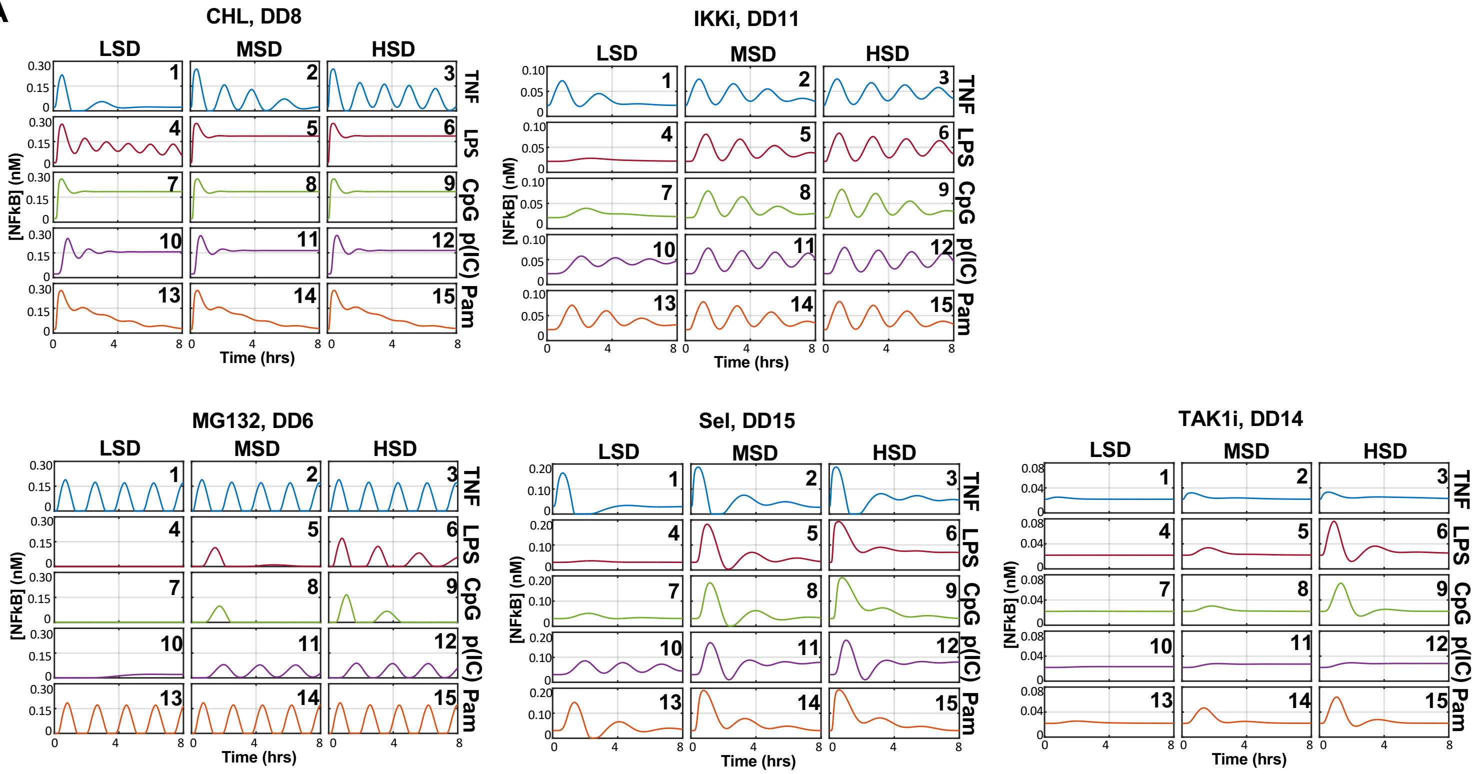


B

| Drug regime | CHL, DD8 | IKKi, DD11 | MG132, DD6 | Sel, DD15 | TAK1i, DD14 |
|---|---|---|---|---|---|
| Objective clusters | [1],[2],[3],[4], [5,6,7,8,9], [10],[11,12], [13,14,15] | [1], [2,5,9,14,15], [3,6],[4,7], [8,13],[10], [11,12] | [1-3,13-15], [4,7,10],[5,8], [6],[9],[11,12] | [1],[2],[3],[4], [5,8],[6],[7], [9,14,15],[10], [11,12],[13] | [1-5,7,8, 10-14], [6],[9,15] |

C

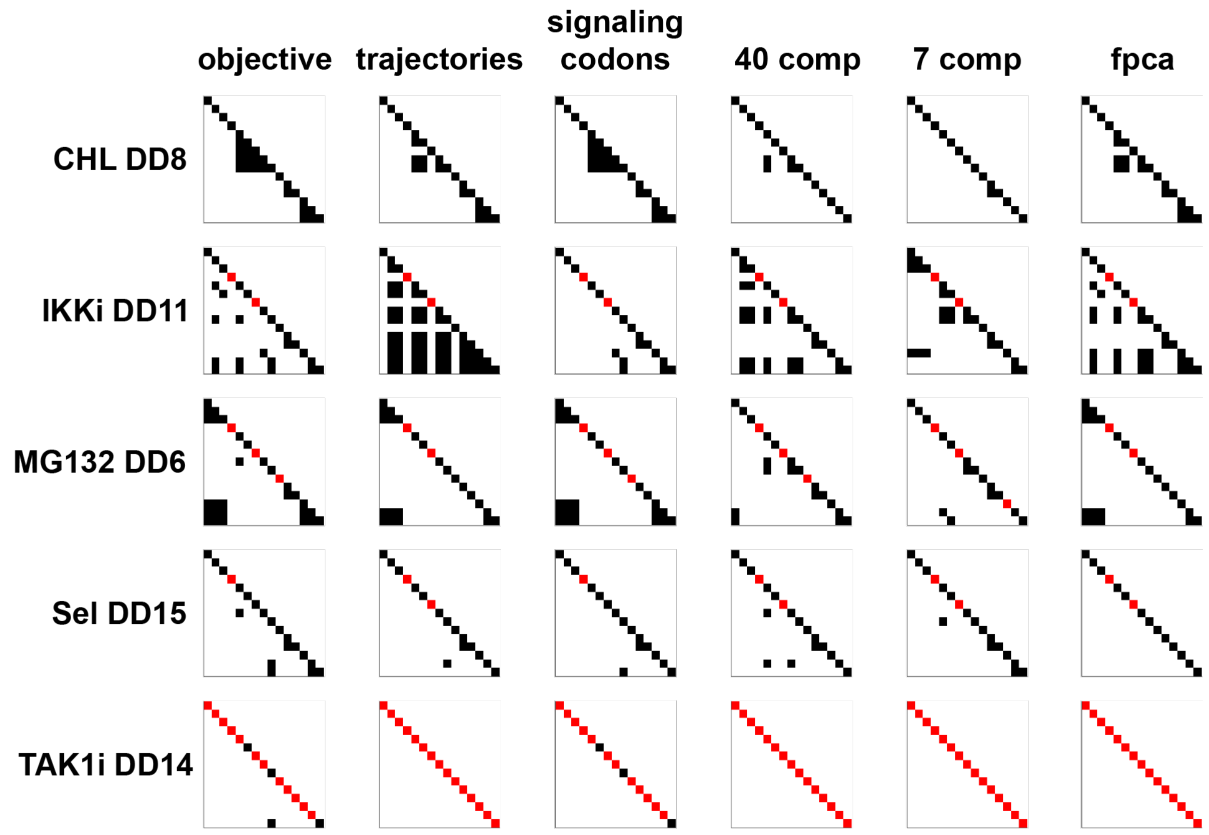

**Figure S3 Construction of stimulus cluster maps for selected drug regimes**

**(A)** Trajectories of nuclear NFκB concentration (y-axis of subpanels) over time (x-axis of subpanels) for drug regimes CHL DD8, IKKi DD11, MG132 DD6, Sel DD15, TAK1i DD14, and across 5 ligands and 3 doses. Solid colored lines represent the trajectories under drug treatment, while dashed lines depict untreated trajectories. Rows indicate different ligands (labeled on the right of Figure 3D panel), and columns specify ligand doses (labeled on the top of the panel). Stimulation indices are labeled in the top right corner.

**(B)** Objective clusters for 5 representative regimes: CHL DD8, IKKi DD11, MG132 DD6, Sel DD15, and TAK1i DD14 used in epsilon network clustering. Clusters for each regime are annotated according to the trajectory indices in (A).

**(C)** Stimulus cluster maps constructed from epsilon network clustering results for 5 representative regimes. Each row and column within one map corresponds to a specific stimulus, as denoted on the left side of the left panels. Within each map, off-diagonal black squares represent responsive clusters and red squares on the diagonal represent inhibited NFkB signaling (non-responder). Panels from left to right display the objective clusters, clusters derived from the trajectory space, signaling codon space, 7 component CPD feature space, and 40 component CPD feature space, and fPCA feature space (Labeled on the top of the panel).

### Figure S5

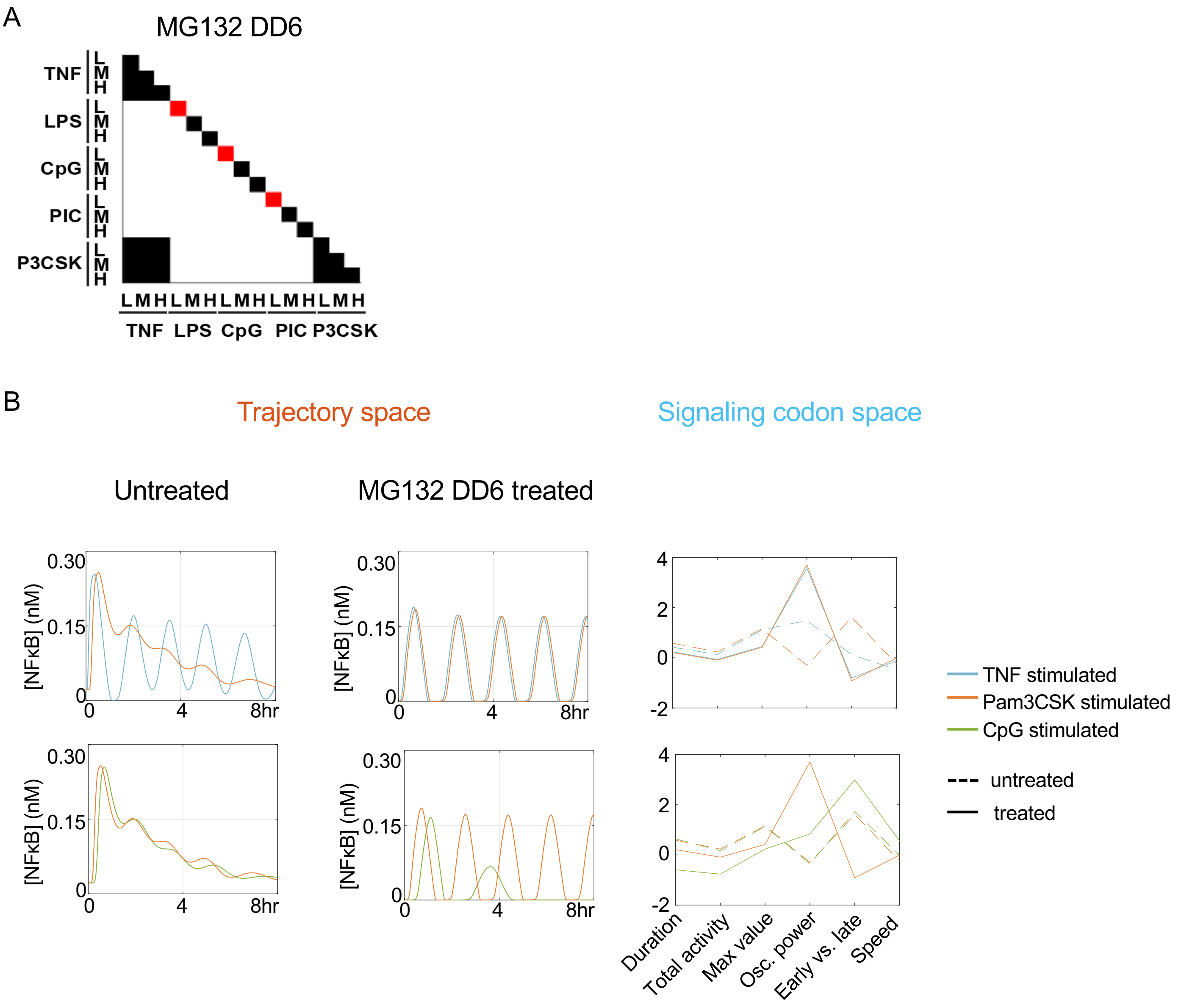


**Figure S5 Decoding stimulus-response confusion, specificity, and inhibition scores for an example drug treatment**

(A) Stimulus cluster maps for trajectories treated with MG132 at DD6.

(B) Examples of decreased temporal coding capacity (high doses of TNF and Pam3CSK, top row) and increased temporal coding capacity (high doses of Pam3CSK and CpG, bottom row), represented in the trajectory space (left and middle columns) and signaling codon space (right column).

### Figure S6

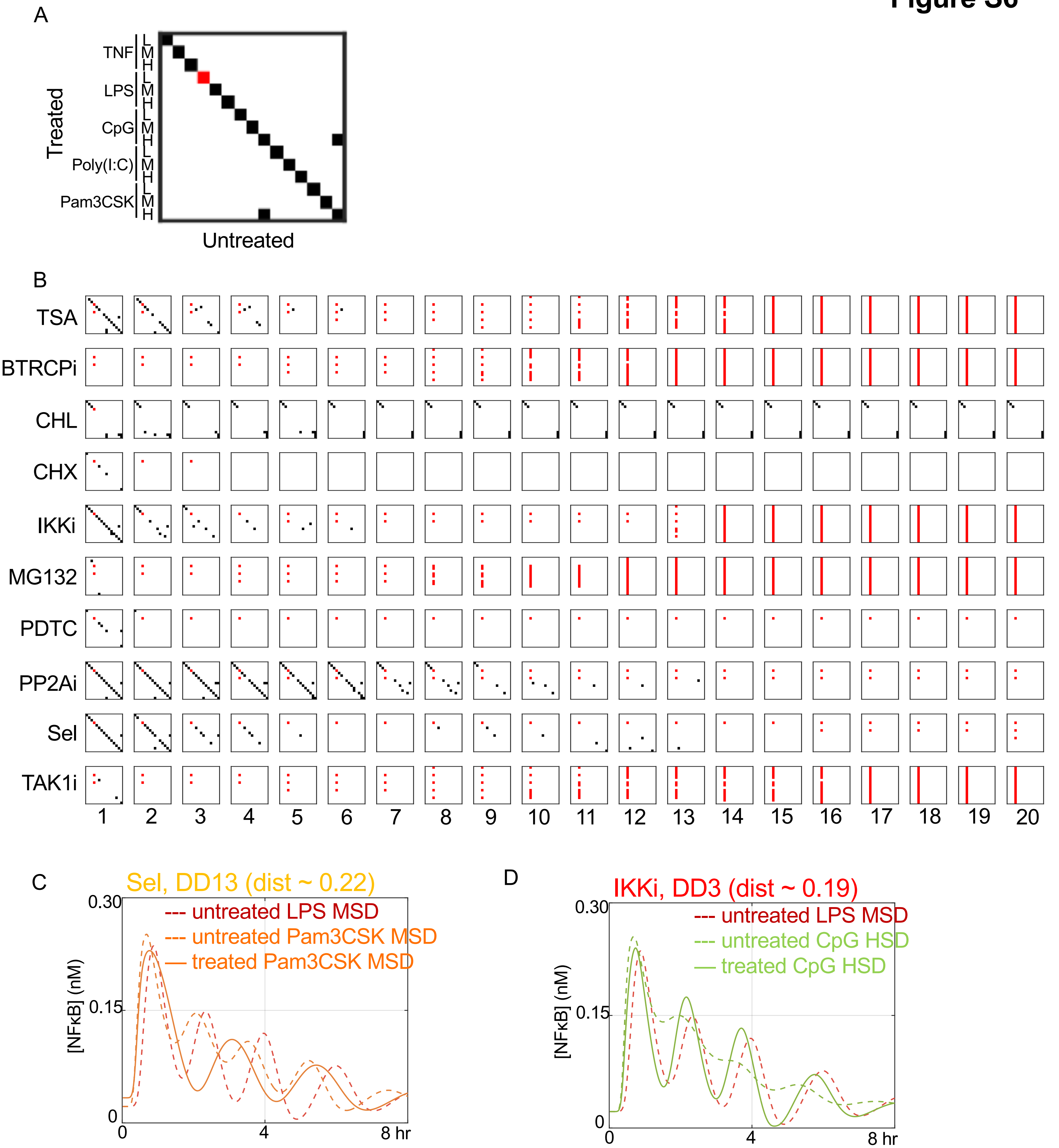


**Figure S6 Signaling-codon derived visualization of temporal coding capacity for 200 drug regimes vs. the 15 stimulated, untreated NFκB responses**

Stimulus cluster map for the 15 stimulus conditions under no drug treatment (untreated) vs. 15 stimulus conditions under (A) an example drug regime and (B) all 200 drug regimes. Columns represent untreated stimuli and rows represent treated stimuli. Black clusters on and off the diagonal represent "confusion" between treated and untreated NFκB responses under the same or different stimulus condition, respectively. and off-diagonal. Red squares represent inhibited NFκB signaling (non-responder). (C-D) Examples of confusion and distinction of the stimuli before and after drug treatment.
